# Single cell transcriptomics delineates the immune-cell landscape in equine lower airways and reveals upregulation of the FKBP5 gene in horses with asthma

**DOI:** 10.1101/2022.09.27.509660

**Authors:** Miia Riihimäki, Kim Fegraeus, Jessica Nordlund, Ida Waern, Sara Wernersson, Srinivas Akula, Lars Hellman, Amanda Raine

## Abstract

**Background:** Equine asthma (EA) is a heterogenous, complex disease with a significant negative impact on horse welfare and performance. EA and human asthma have fundamental similarities, making EA a useful large animal disease model. Bronchoalveolar lavage (BAL) fluid provides a snapshot sample of the immune cells occupying the alveolar space and is one of the most relevant sample types for studies of chronic inflammation in the lung. In this study, we sequenced single equine BAL-cells in order to study the immune cell landscape of the respiratory tract of horses diagnosed with mild-to-moderate EA and healthy controls.

**Results:** ScRNA-seq analysis of ~63,000 cells from eleven horses diagnosed with mild-moderate asthma (mEA) and eight healthy controls were performed using the Drop-Seq technology. We identified five major immune cell populations in equine BAL; alveolar macrophages (AM), T cells, neutrophils, mast cells and dendritic cells, as well as subtypes thereof. The cellular subtypes demonstrated herein have previously not been characterized in horses. Differential gene expression analysis revealed upregulation of genes in mEA horses, including FKBP5 and CCL24, which have previously been associated with asthma in other species. The most significantly upregulated gene across the cell types in EA was FKBP5, a chaperone protein involved in regulating the assembly, activity, and sensitivity of the glucocorticoid receptor

**Conclusion:** Herein we demonstrate the first comprehensive scRNA-seq map of the immune-cell populations in BAL from horses with asthma and healthy individuals. The glucocorticoid receptor associated protein FKBP5 was identified as a potential biomarker for EA.

## Introduction

Asthma is one of the most prevalent chronic diseases affecting the airways of humans and certain domestic animals, including horses. More than 330 million people worldwide are affected by asthma, and in 2019 it caused more than 450,000 deaths worldwide, making this disease a major health burden (WHO). Traditionally, mouse models are used to study the underlying mechanisms of asthma. However, unlike humans, mice do not spontaneously develop asthma and the disease must be artificially induced, for example by strong allergens (1). In contrast, horses are one of few animal species that spontaneously develop asthma (2). Equine asthma (EA) is a major horse welfare problem, and it is one of the dominant causes of poor performance in athletic horses. Like its human equivalent, EA is a heterogeneous, complex disease associated with both genetic and environmental factors. Two of the main environmental factors influencing the prevalence and severity of the disease are stable conditions and feed quality (3,4), which are both related to the exposure to dust and mold. However, EA has a complex genetic background (4–10). The clinical symptoms of EA are similar to human asthma and include coughing, airway obstruction, increased respiratory effort and mucus accumulation, as well as decreased athletic performance (11–14). EA is currently subdivided into severe equine asthma (sEA, previously known as recurrent airway obstruction, RAO) and mild-moderate equine asthma (mEA, previously known as inflammatory airway disease, IAD) (11,15). The categorization into a severe and mild form of the disease is used to define cases clinically and is, apart from severity of symptoms, also based on the presence and frequency of certain inflammatory cell types in bronchoalveolar lavage (BAL) (16). However, the two conditions sometimes share clinical, cytological as well as functional similarities and at present there are no definite criteria or biomarkers which can be used to clearly distinguish between mEA and sEA (16). It is likely that additional endotypes exists under the EA umbrella term and that the division into mEA and sEA in horses is too coarse (17). Nonetheless, given the similarities between EA and human asthma (15,18), the horse presents an excellent model for studying the pathological processes of disease.

EA is traditionally diagnosed based on medical history, clinical examination, airway endoscopy findings, lung-function tests, and BAL cytology (16). Studies have demonstrated a shift in cell composition (increased granulocyte infiltration) in the lungs of horses as well as humans diagnosed with asthma compared to healthy controls, making the BAL cytology a valuable tool for diagnosing the disease (19,20). Traditional BAL cytology, however, does not provide sufficient resolution to study disease mechanisms or perform high resolution sub-classification of asthma. High throughput sequencing can be valuable to obtain more detailed information based on gene expression and have been used in several studies to study gene expression differences in BAL-cells and bronchial biopsies sampled from horses diagnosed with asthma (7,21–24). Traditional bulk RNA-sequencing technologies obtains measurements from a mixture of cells in which inter-cellular differences are averaged. In contrast, single cell RNA sequencing (scRNA-seq) is a technology that offers possibilities to analyze gene expression at the individual cell level, which enables characterization of cellular diversity. Although many studies have been performed with cells of human and mice origin, there are very few studies published using scRNA-seq on horse. To our knowledge there is yet no comprehensive cell atlas of the immune cell landscape in the horse respiratory tract (25,26). In this study, we provide novel and detailed information of the cell types occupying the alveolar space of horses in health and disease, by performing sRNA-seq on clinically relevant BAL samples and controls. Moreover, we identify significantly differentially expressed (DE) gene transcripts of relevance for EA, which can be further explored as potential biomarker for the disease and for prediction of therapy response in horse and human.

## Results

### The global cellular landscape in the lower airways of horses in health and disease as detected by scRNA-seq

Cell compositions in the BAL samples from 19 horses (11 cases with mEA, 8 controls) were assessed by cytology staining during routine diagnostic procedures (Table 1). The neutrophil proportions in the BAL samples were in the normal to slightly elevated range (1-7 % in both the asthma & control groups), except for three horses who presented BAL neutrophil levels > 10% (two EA and one control). Strikingly, ten out of the eleven horses in the EA group had elevated mast cell counts compared to the healthy controls (4-11 % vs 1-2 %, p < 10^−4^Students t-test) (Table 1). Moreover, six of the horses in the EA group had elevated eosinophil counts compared to the control group (3-18 % vs 0-1 %, p = 0.05) (Table 1). The healthy control horses had normal cell counts in BAL, except for the one horse that had slightly elevated neutrophil counts. One of the control horses had elevated total number of leucocytes in BAL (horse H, table 1) and had a history of occasional single coughs. Horse FN, Table 1, had a normal BAL cytology at the sampling occasion, despite clinical history of seasonal mEA.

**Table 1.** Total BAL cell counts, BAL cytology data and number of cells analyzed with DropSeq per horse.

| ID | Pheno-<br>type | Leukocytes<br>(10e6/L) | Neutrophils<br>(%) | Lymphocytes<br>(%) | Macrophages<br>(%) | Eosinophils<br>(%) | Mast cells<br>(%) | No of<br>cells* |
| --- | --- | --- | --- | --- | --- | --- | --- | --- |
| MW | Asthma | 320 | 16 | 48 | 28 | 3 | 5 | 3367 |
| O | Asthma | 400 | 4 | 22 | 49 | 18 | 7 | 5224 |
| Z | Asthma | 550 | 18 | 28 | 45 | 5 | 4 | 2716 |
| N | Asthma | 310 | 0 | 37 | 47 | 6 | 10 | 3421 |
| FS | Asthma | 190 | 7 | 33 | 42 | 8 | 10 | 3003 |
| C | Asthma | 460 | 2 | 41 | 50 | 0 | 7 | 2709 |
| VE | Asthma | <100 | 3 | 37 | 52 | 1 | 7 | 4461 |
| VA | Asthma | 280 | 5 | 12 | 71 | 6 | 6 | 3060 |
| T | Asthma | 190 | 5 | 33 | 53 | 0 | 9 | 3393 |
| D | Asthma | 180 | 1 | 35 | 52 | 1 | 11 | 3092 |
| FN | Asthma | 140 | 1 | 36 | 61 | 0 | 2 | 2067 |
| MY | Control | 350 | 3 | 46 | 50 | 0 | 1 | 3885 |
| A | Control | 250 | 7 | 30 | 62 | 0 | 1 | 2732 |
| B | Control | 470 | 13 | 36 | 49 | 1 | 1 | 3326 |
| F | Control | 250 | 3 | 44 | 51 | 0 | 2 | 3292 |
| H | Control | 510 | 5 | 44 | 50 | 0 | 1 | 3625 |
| Q | Control | 150 | 6 | 29 | 64 | 0 | 1 | 3437 |
| G | Control | 200 | 3 | 31 | 63 | 1 | 2 | 4207 |
| P | Control | 170 | 4 | 30 | 65 | 0 | 1 | 3705 |

\*Nr of single cell transcriptomes recovered per horse after quality filtering

Single cell RNA sequencing of cells in BAL from the 19 horses included in the study were performed using the Drop-Seq method (study outline in Figure 1A) (27). After single cell encapsulation, sequencing and filtering of data using stringent quality thresholds (see Methods and Supplementary Figure 1), the single cell transcriptomes from in total 63,022 cells remained for integrated clustering and cell type annotation (asthma n=36,513 cells, healthy n= 28,209 cells). In total, 11,844 genes were detected.

**Figure 1.**
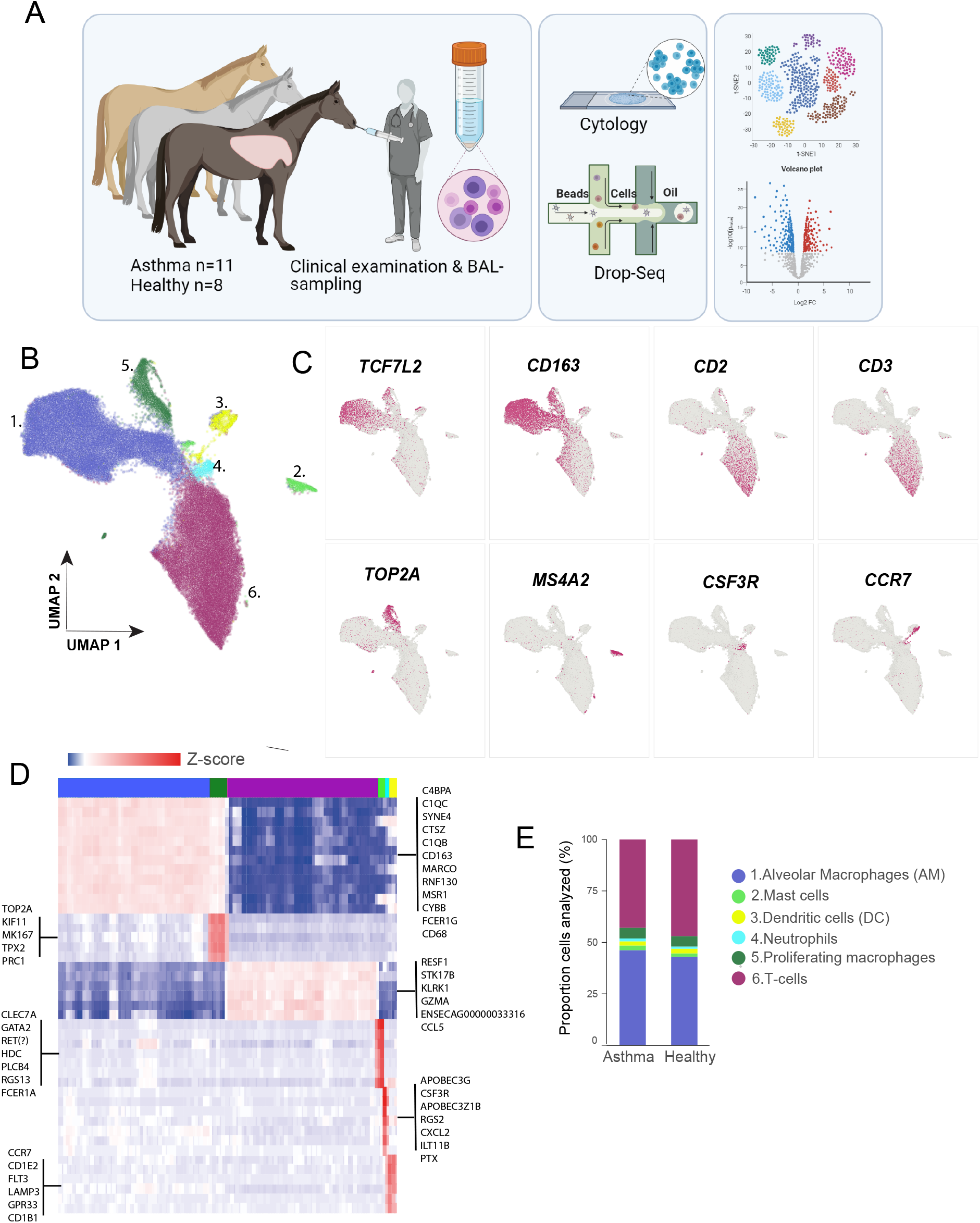
Initial clustering of BAL cells. A) Graphical illustration of the study design. B) UMAP representation of 63,022 equine BAL cells colored by the six major cell-groups; macrophages (28,277 cells, #1) proliferating macrophages (3420 cells, #2), T-cells (28,112 cells, #3), mast cells (1257 cells, #4), neutrophils (797 cells, #5) and dendritic cells (1339 cells, #6. C) Example of expression of specific markers for the different cell types. D) Heatmap demonstrating the difference in expression levels of the top biomarker genes for each of the six major clusters. E) Proportion of cell-types in the mEA and healthy control horses by group.

Clustering of the major cell populations in BAL were visualized by UMAP (Figure 1B) (28). As expected, the most abundant immune cell types were alveolar macrophages (AMs) and T cells. AMs were identified based on the expression of established canonical macrophage markers: *MARCO, CD163, CD68*, and *TCF7L2* (Figure 1C and D and Supplementary File 1). We identified a substantial population (3,077 cells) of proliferating AMs which, in addition to canonical macrophage markers genes, also specifically expressed differentiation and cell division genes, such as *TOP2A, KIF11, MK167* and *TPX2*. T cells were annotated based on expression of classical surface receptor T cell markers (*CD*2, *CD3*), cytotoxicity genes (*GZMA, KLRK1*), chemokine (*CCL5*), T cell receptor (*TRCB1/ENSECAG00000033316*), and the serine/threonine kinase *STK17B* (Figure 1C and D). We also found a sizable subgroup of cells (n=2,800) that clustered with T cells, but additionally co-expressed the canonical macrophage markers (double positive cells) (Supplementary Figure 1). T cell/monocyte double positive (DP) cells have been observed previously e.g., in blood (29,30). It is likely that the DP cells observed herein represent either technical doublets (where two or more cells enter the same droplet by chance) and/or captures functional T cell/AM complexes in the same droplet. The DP cells were excluded from the downstream analysis.

Mast cells were annotated according to expression of tryptase *TPSB2*, the high affinity IgE receptor beta and alpha chains (*MS4A2, FCER1A*) and the transcription factor *GATA2*. A small cluster of neutrophils (Figure 1B, cluster 4) were annotated according to expression of the chemokine *CXCL2*, the granulocyte colony stimulating factor *CSF3R, APOBEC*-family transcripts and *CD85 (ILT11B*). Another small cluster (Figure 1B, cluster 3) were characterized by high expression of *CD1, CCR7, LAMP3* and *FLT3*, which are known markers for dendritic cells.

When comparing the scRNA-seq data to the routine cytology analysis, the majority of cells detected were AMs and T cells, as expected (Figure 1, Table1). The proportions of granulocytes were lower in the scRNA-seq data compared to the cytology analysis. Notably, our clustering did not confidently designate an eosinophil cluster, despite the elevated eosinophil BAL cytology counts observed for a subset of the EA horses included in the study. Although there was a clear difference in mast cell counts (from cytology) between the EA and healthy controls (Table 1), comparison of the proportions of the various cell types between the asthma and healthy group as identified by scRNA-seq showed no major difference between the groups (Figure 1E, Supplementary Figure 2A, Supplementary File 9).

### Distinct subgroups of equine T cells populate the alveolar compartment of horses

In order to increase the cluster resolution and to facilitate annotation of different T cell populations, we performed independent sub-clustering of 23,480 T cells, excluding the putative T cell-macrophage DP cells. This initially resulted in six T cell clusters (Supplementary Figure 3), where four contained clear signatures based on previously defined T cell phenotype (Figure 2A and B, Supplementary Figure 3). Focusing on the genes expressed in those four T cell clusters, two clusters were annotated as CD8^+^ T cells (hereafter denoted CD8^+^_TR_ and CD8^+^_EM_), one cluster as CD4^+^ T cells, and one cluster as γδ-T cells. The four major T cell types were found in all horses, and we observed no significant difference in the proportions between the two conditions (Figure 2C, Supplementary Figure 2B, p-values are found in Supplementary File 9).

**Figure 2.**
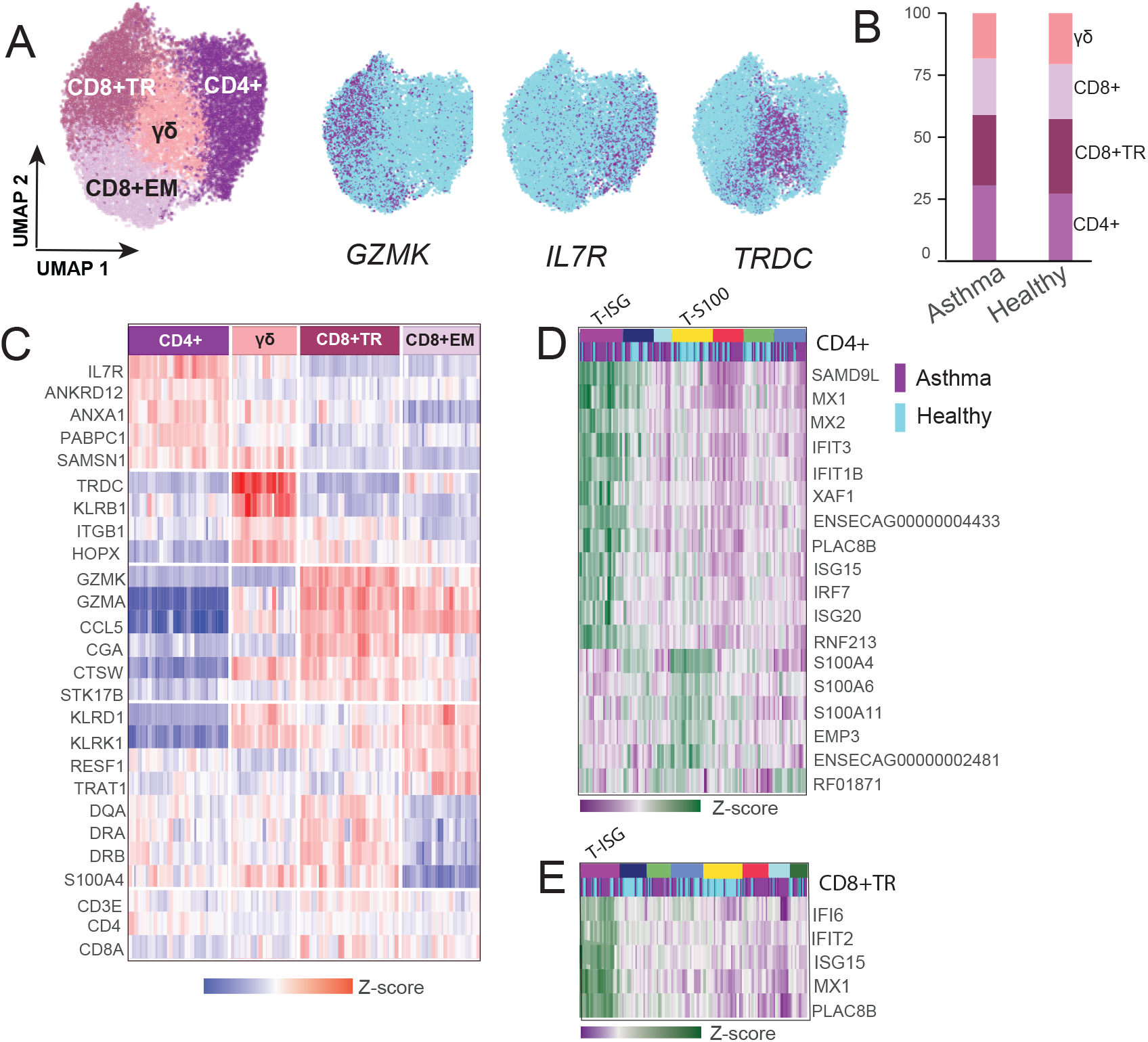
Re-clustering of T cells. A) UMAP plot illustrating the four equine BAL T cell groups. Expression of top differentially expressed markers mapped onto the UMAP-plot are shown to the right. B) Differential expression of T cell cluster markers visualized with a heatmap. C) Proportion of T cell subtypes in the asthma vs healthy group of horses. D) Iterative clustering shows seven subpopulations of CD4^+^ cells. The purple subset (T-ISG) expresses interferon stimulated genes and the yellow subset (T-S100) expresses genes involved in cytoskeleton organization and cell adhesion. The cells from EA horses are shown in purple and the controls are shown in blue. E) Iterative clustering of the CD8^+^TR cluster shows eight subclusters. The purple subset (T-ISG) selectively expresses interferon stimulated genes.

The cells in the CD8^+^ clusters expressed genes associated with cytotoxicity as well as known CD8^+^ T cell markers (Figure 2A and B). We identified 48 DE genes (FC >2) between the two CD8^+^ T cell clusters (Figure 2B, Supplementary Figure 3, Supplementary File 2). Gene ontology (GO) pathway analysis using the set of genes with > 2-fold higher expression in the CD8^+^_TR_ cluster indicated enrichment to biological pathways involved in adhesion, migration, and T cell activation (Supplementary Figure 3B). The CD8^+^_TR_ cluster expressed higher levels of transcripts typical of cytotoxic CD8^+^ T cells, such as granzymes (*GZMA, GZMK*), the cytotoxicity associated cathepsin W (*CTSW*), killer lectin (*KLRK1*) and chemokines (*CCL5*). In the same cluster we also detected higher transcription (compared to the CD8^+^_EM_ cluster) of integrins and other genes implicated in adhesion and migration (*IGTAE, ITGB, S100A4, S100A11* and *ANXA1*). The expression of MHC class II genes (*DQA, DQB, DRA* and *DRB*) was also elevated in the CD8^+^TR cluster. Interestingly, we identified higher expression of the *CGA* gene (the alpha chain of glycoprotein hormones) in this cluster. To our knowledge, this gene has not previously been described in T cell function. Taken together, the expression signature of the CD8^+^_TR_ cluster suggests a population of active tissue resident CD8^+^ memory cells.

The CD8^+^_EM_ cluster was also characterized by high *GZMA, GZMK* and *CCL5* expression, albeit lower *GZMK* expression than the CD8^+^_TR_ cluster. Moreover, expression of *KLRD1, RESF1* and *TRAT1* were higher in this cluster (Figure 2 and Supplementary Figure 3C). The lower levels of CD103 (*ITGAE*) and other transcripts related to adhesion in this cluster indicate a population of circulating effector memory T cells (CD8^+^_EM_) (31).

In comparison to the other T cell clusters, the CD4^+^ cluster was characterized by the upregulation of *IL7R, ANXA1, PABPC1, TNFSF13B* and *SAMSN1*. A distinct subset of CD4^+^ cells displayed higher expression of S100 family genes (*S100A4, S100A6* and *S100A11*)as well *EMP3* (Epithelial membrane protein 3) a protein believed to be involved in cell proliferation and cell-cell interactions, plus other genes involved in cytoskeleton organization (Supplementary File 2, Supplementary Figure 5). Notably, this cluster was found to mostly comprise T cells derived from the healthy control horses (p-value =0.002. Figure 2D and Supplementary Figure 2D). We were, however, not able to subclassify the CD4^+^ cluster into any of the conventional Th-subtypes (Th1,2,17, T-reg) based on the single cell expression signatures obtained in this study.

Of note, we identified cells within the CD4^+^ and CD8^+^_TR_ clusters with significant upregulation of interferon stimulated genes (ISG), including *MX1, PLAC8D, IFIT3*, and *ISG15* (Figure 2D, Supplementary Figure 4, Supplementary File 2). Upon further iterative clustering of the CD4^+^ and CD8^+^_TRM_ populations, the ISG expressing T cells (T-ISG^Hi^) fell into distinct sub-clusters (Figure 2D). While the T-ISG^Hi^ cells were found in the majority of horses, a trend was observed towards higher proportions of these cells in EA (FC= 2.8, p-value = 0.10 in the CD4^+^ cells) (Supplementary Figure 2D). Two horses (N and C) in the EA group exhibited a stronger ISG expression signature in both T cells an AMs compared to all other horses (Supplementary Figure 5).

The γδ T cells were characterized by high T cell receptor delta constant *TRDC*, lower alpha constant *TRAC* and high *CTSW*. Moreover, the γδ-T cell population exhibited a high level of transcripts for the killer lectin receptor *KLRB1* and also expressed *KLRD1* and *KLRK1*. Altogether, this suggest cytotoxic potential and is in agreement with what was demonstrated for circulating γδ-T cells in a previous scRNA-seq study of the equine PBMC fraction (26).

Although the two remaining clusters (T_NA_ and T_DP_) expressed canonical T cell markers, they did not express markers that could enable further sub-annotation to a specific T cell subtype (Supplementary Figure 3, Supplementary File 2).

By performing iterative clustering on a large population of T cells we thus outlined the major subtypes present in the equine respiratory tract and also demonstrate novel T-ISG^Hi^ and T-S100^Hi^ equine T cell populations.

### Heterogeneity among equine alveolar macrophages

In a similar manner as for the T cells, we performed re-clustering of AMs (excluding the cluster of proliferating AMs). As AMs are large cells with relatively high RNA content, we applied the stringent cut-off of including only cells expressing > 500 genes. This resulted in 22,000 AM cells for downstream iterative clustering (Figure 3A). Given that recent studies have demonstrated considerable heterogeneity among human and mouse AMs, high diversity is to be expected also for horse AMs (32,33). Sub-clustering revealed several equine AM subpopulations with distinct gene expression profiles (Figure 3A). As for the T cell sub-populations, we did not find a pronounced difference in the proportions when comparing EA and healthy controls (Figure 3C, Supplementary Figure 2C, Supplementary File 9). Within the AMs, the cluster denoted AM4 had the largest number of expression differences compared to the other clusters (>300 differentially expressed genes compared to cluster AM3, Supplementary Figure 5 and Supplementary File 3). Cluster AM4 was characterized by higher expression of the glycoprotein Nmb *GPNMB*, galectins; *LGALS3, LGALS1* and members of the S100 gene family (*S100A4, S100A6* and *S100A11*). Other upregulated genes included superoxide dismutase *SOD2*, phospholipase *PLD3*, cathepsins (*CTSB, CTSD*) and the C15orf48 orthologue; *ENSECAG00000012148*. The macrophage scavenger receptor (*MARCO*) transcript levels were notably lower in AM4 compared to all the other macrophage clusters (Figure 3B). Of note, human monocyte-derived AMs are previously shown to express lower levels of MARCO compared to embryonically derived AMs (34).

**Figure 3.**
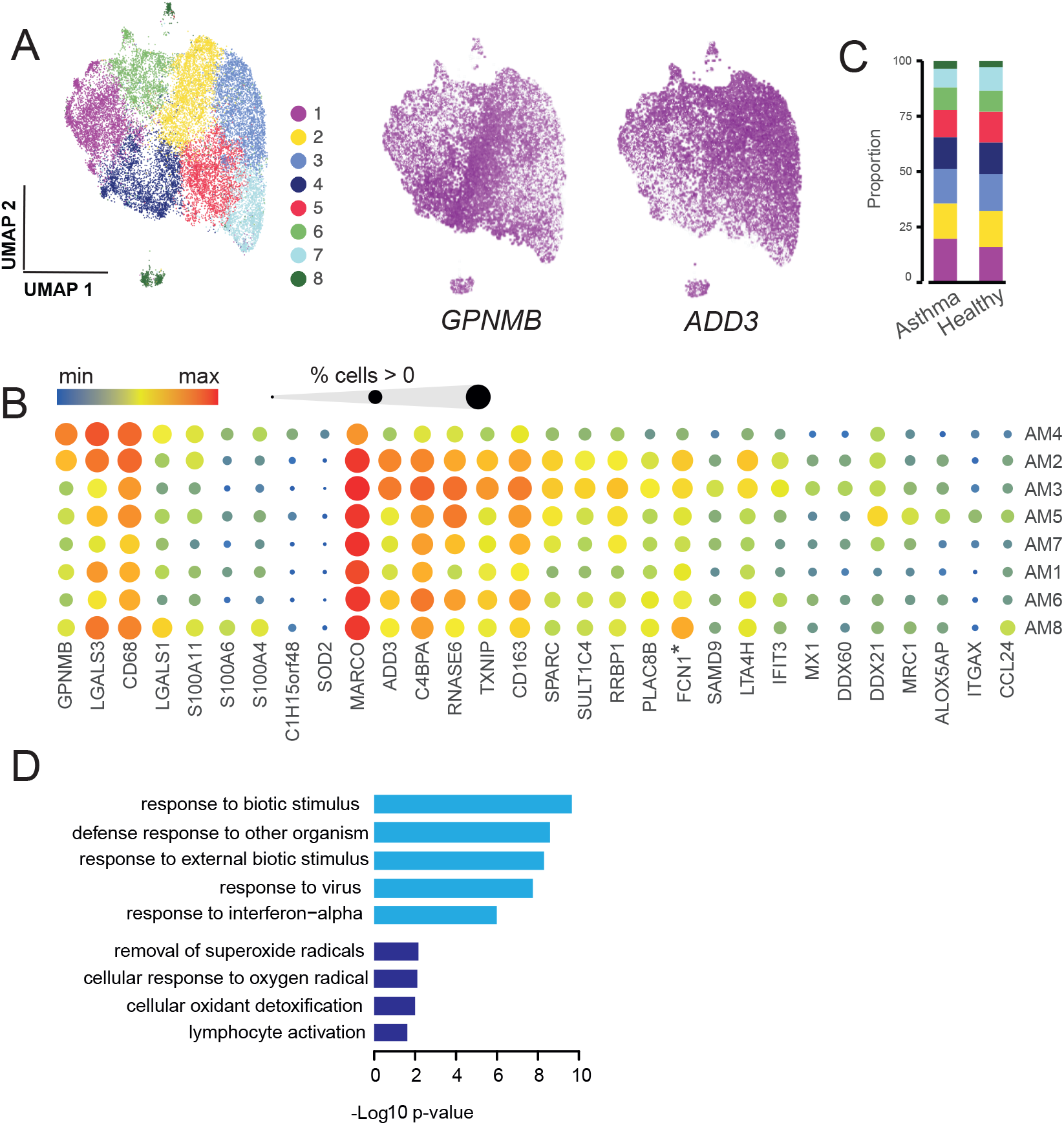
Reclustering of alveolar macrophages. A) UMAP illustration of eight alveolar macrophage (AM) subsets. The top differentially expressed markers for cluster 4 (AM4) and cluster 3 (AM3) were glycoprotein Nmb (*GPNMB*) and γ-adductin (*ADD3*), respectively. B) Bubble plot visualizing the expression levels for a number of cluster marker genes. C) Proportion of AM clusters in the AE and healthy groups. D) Genes upregulated in cluster AM3 (cluster 3) were significantly enriched in the GO-pathways colored in light blue. Genes upregulated in AM4 (cluster 4) were significantly enriched in GO-pathways colored in dark blue. *\*FCN1= ENSECAG00000000436*

Markers traditionally used to identify pro-inflammatory M1-polarized (*CD68*) and anti-inflammatory M2-polarized macrophages (*CD206, CD163*) respectively, were expressed across all the cellular AM subtypes. This observation confirms the emerging view that the classical M1/M2 polarization is an inappropriate classification of AMs. *CD68* expression was moderately higher in the AM4 cluster and concurred with lower *CD206* and *CD163* expression. The reverse was observed in AM3 and AM5 clusters (Figure 3B). Cluster AM3 exhibited a significant ISG transcript signature (i.e., increased expression of *SAMD9L, MX1, IFIT2/3, XAF1, DDX60* etc., Figure 3B and Supplementary Figure 6), which is not in accordance with a classical anti-inflammatory M2 phenotype. The most significantly upregulated gene in the AM3 cluster was the membrane skeletal protein *ADD3 f* Adductin-3).Other highly expressed genes were the thioredoxin interacting protein *TXNIP, RNASE6* as well as complement-system genes Ficolin-1 (*FCN1|ENSECAG00000000436*) and *C4BPA*.GO pathway analysis based on the genes upregulated in the AM3 cluster showed enrichment in pathways involved in response to viruses, other organisms, external biotic stimulus and interferon-alpha. In contrast, genes upregulated in AM4 were enriched in pathways related to oxidant detoxification, removal of superoxide radicals and lymphocyte activation (Figure 3D).

Notably, the cells in cluster denoted AM2 were characterized by an expression pattern comprising elements of both the AM3 and AM4 clusters. Thus, the AM2 cluster cells expressed both higher *GPNMB, LGALS4* and *CD68* compared to AM3, as well as higher *MARCO, ADD3* and *CD163*. Moreover, expression of interferon stimulated genes was found within the AM2 cluster (Figure 3D). The cluster denoted AM5 were mainly characterized by (compared to AM3 and AM4) lower levels of *GPNMB, LGALS3, ADD3* and ISG transcripts as well as higher expression of e.g., *MRC1* (CD206), *CD163, ITGAX (CD11c), DDX21, ALOX5AP* and *ENSECAG00000039383* (novel gene, *Wfdc21* orthologue) (Figure 3B, Supplementary File 3). The remaining AM clusters (AM1, AM6 and AM7) were rather characterized by overall lower number of genes detected (Supplementary Figure 6). The small cluster of AM8 were characterized by higher expression of ribosomal protein mRNAs compared to the other AMs.

### Equine BAL mast cell and neutrophil populations

Next, we examined the BAL mast cell and neutrophil populations. Although the granulocyte recovery was lower compared to the numbers expected from BAL cytology analysis, the single cell transcriptomes of ~1,250 mast cells (831 cells from mEA, 426 cells from controls) and ~800 neutrophils were recovered. In addition to the markers used for cluster identification shown in Figure 1, the mast cells in equine BAL expressed high levels of mRNAs in histamine, prostaglandin and leukotriene production pathways (*HDC, HPGDS, PTGS1* and *LTC4S*). Furthermore, we detected expression of mast cell specific genes such as *PLCB4, RGS13* and *SIGLEC6* (Figure 1, Supplementary File 1). Whereas the tryptase *TPSB2* was highly expressed in the mast cells, no expression of chymase (*CMA*) or carboxypeptidase A3 (*CPA3*) was detected. Taken together, this reveals that the mast cells in the lower airways of the equine lung primarily are of the classical mucosal subtype (35).

Re-clustering of the mast cells resulted in six subpopulations. The mast cells clusters were rather similar, although the cells in clusters 1 and 4 had the most prominent expression profiles (Figure 4 and Supplementary File 4). Cluster 1 nearly exclusively comprised cells from the EA group. This cluster expressed higher levels of leukotriene C4 synthase (*LTC4S*)as well as the glucocorticoid receptor associated peptidyl-prolyl cis-trans isomerase *FKBP5*. Moreover, regulator of G-protein signaling 1 (*RGS1*), a gene which is associated with auto-immune disorders and asthma in humans, was higher expressed in cluster 1. In cluster 4 we observed complement C1q transcripts (*C1QB* and *C1QC), CD74, IFI27, MNDA* and other genes which are also expressed in macrophages (Figure 4A).

**Figure 4.**
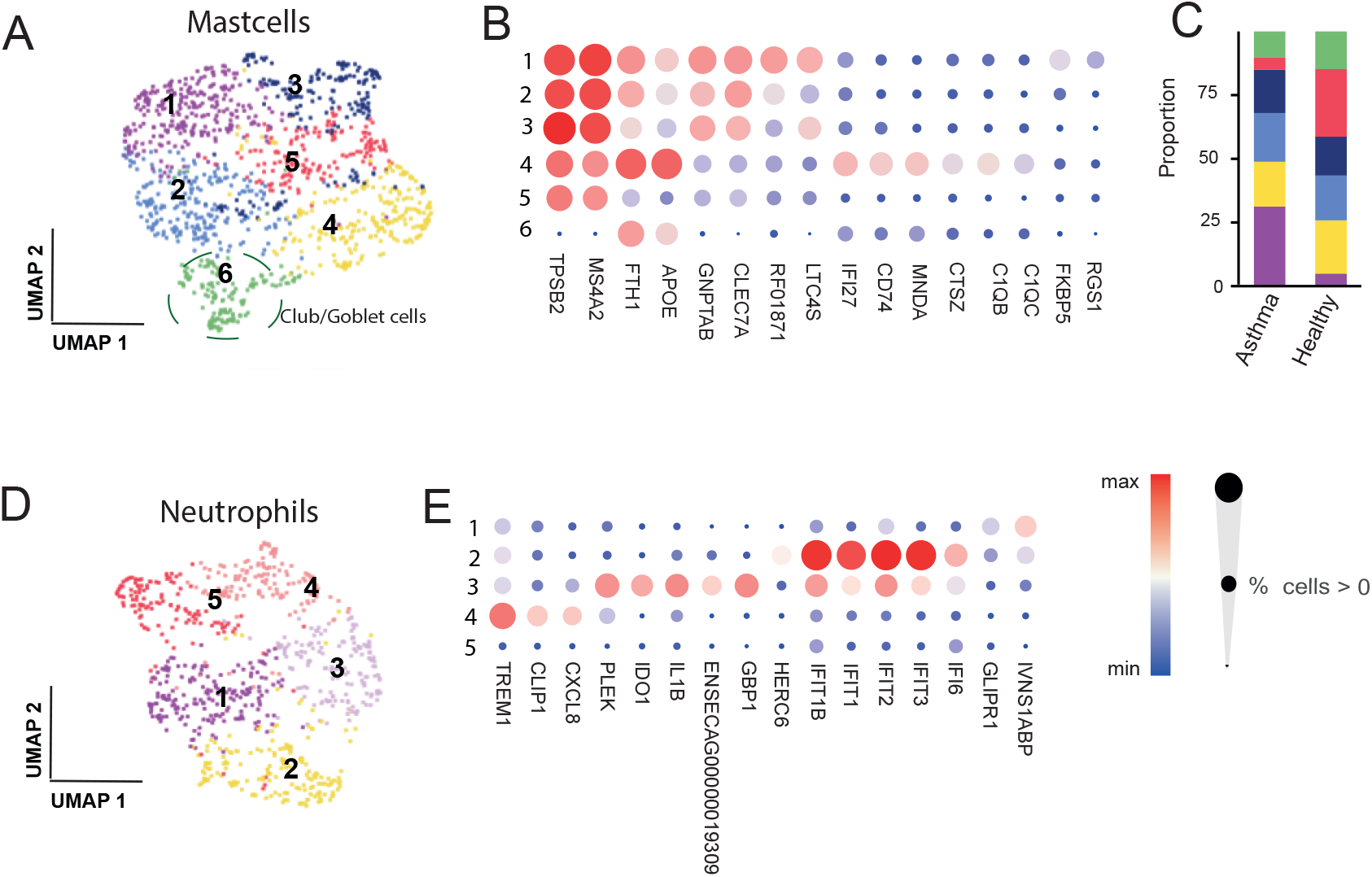
Reclustering of mast cells and neutrophils. A) UMAP illustration of mast cell clusters. B) Bubble plot visualizing the expression levels of the cluster marker genes. C) Proportions of mast cell subpopulations in asthma and healthy horses. D) UMAP illustration of five neutrophils subclusters. E) Bubble plot visualizing the expression levels of the neutrophil cluster marker genes.

A small population (<100 cells), clustered closely together with the mast cells although absent of typical mast cell signatures. A marker signature including *CLCA1, VMO1, TFF3, SCGB1A1* and *CXCL17* suggests that these cells are club or goblet cells derived from the airway epithelium (Figure 4A, Supplementary File 4).

The BAL neutrophils had overall lower gene expression as is expected from a terminally differentiated granulocyte population. Even though the total number of neutrophils analyzed herein were rather low (n=800), we observed several distinct clusters among them. For instance, neutrophil cluster 2 had strong upregulation of ISG expression, while cluster 3 exhibited elevated transcript levels for pro-inflammatory cytokine IL-1B, together with the tryptophan-catabolizing and immunosuppressive indolamine-2, 3-dioxygenase enzyme *IDO1* and pleckstrin (*PLEK*). A third subset of cells (cluster 4) displayed increased expression of the immunoglobulin (Ig) superfamily transmembrane protein *TREM1*, cytokine *CXCL8* (IL-8) and microtubule associated protein *CLIP1* (Figure 4B, Supplementary File 4).

In summary, we observed several subpopulations of granulocytes in the equine lower airways.

### Dendritic cells in equine BAL

The initial clustering of BAL cells singled out a small population of cells expressing dendritic cell markers (n = 1,340 cells, cluster 3 in Figure 1B). As dendritic cell populations in the equine lung have not previously been characterized and little is known regarding the subtypes of dendritic cells populating the equine alveolar compartment, we set out to characterize subpopulations based on transcriptome profiles. Independent reiterative clustering of the dendritic cells revealed six putative subpopulations (Figure 5, Supplementary File 4). Type-2 conventional dendritic cells (cDC2s) were marked by expression of *FCER1A, CD1A, CD1E, CLEC10A* and *CD207*. Furthermore, we designated a cluster of cells expressing high *ENSECAG00000024882* (novel gene, C-C rich chemokine ligand orthologue), *CD14, MS4A7* and *CD68* as monocyte-like cells. Two clusters with distinct expression of *CCR7, FSCN1, LAMP3* and *IDO1* were labelled as activated or migratory dendritic cells (36). A small but distinct cluster of type-1 conventional dendritic cells (cDC1s) exhibited DE genes such as *LRRK2, MT3, CPVL* and *NEXN*. Another small cluster exhibited weak differential expression of T-cell markers *CD3, CXCR6, RORA* and *GATA3*. We were not able to confidently put a label on those cells.

**Figure 5.**
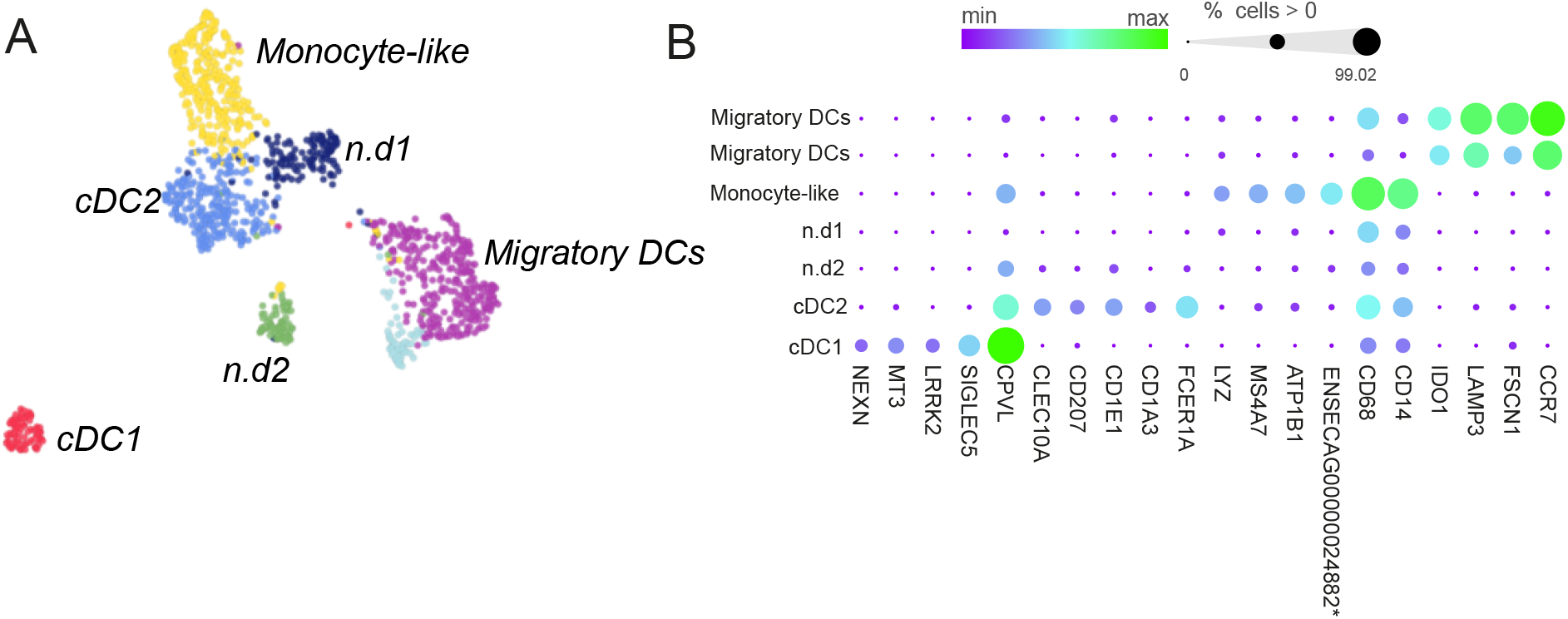
Reclustering of dendritic cells. **A)** UMAP illustration of the dendritic cell clusters. B) Bubble plot visualizing the expression levels of the cluster marker genes. *\*ENSECAG00000024882* unannotated (*CCL15* orthologue). DC =dendritic cells, n.d = not annotated.

### Differential expression analysis reveals FKBP5 and CCL24 as candidate biomarkers for EA

Next, we analyzed DE between EA and healthy controls across different cell types and subpopulations. We performed DE at cell-type level (AMs, T cells and mast cells) using two approaches: i) a pseudo bulk approach with DESeq2) and ii) a two-part Hurdle model equivalent to MAST (37,38). A subset of DE genes is listed in Supplementary Table 1 and the full sets of DE genes are found in Supplementary Files 5,6 and 7.

Interestingly, both approaches identified *FKBP5* as the most differentially expressed gene and the highest fold changes was detected in mast cells (Figure 6, Supplementary File 7). The eosinophil chemotactic protein 2 (eotaxin-2, *CCL24*), was also among the most significant DE genes and highly expressed in AMs derived from EA horses. Whilst *FKBP5* expression was upregulated in all the mEA horses, *CCL24* expression was primarily upregulated in cells derived from four horses (IDs: O, VA, VE, and T) and the highest upregulation was observed in the AM cluster denoted AM5 (Figure 6, Supplementary Figure 7, Supplementary File 5). This finding did, however, not match with individual eosinophil levels in BAL (as measured by cytology, Table 1) and *CCL24* was not among the differentially expressed genes upon comparing mEA horses with elevated eosinophils compared to those with normal eosinophil counts by cytology (data not shown).

**Figure 6.**
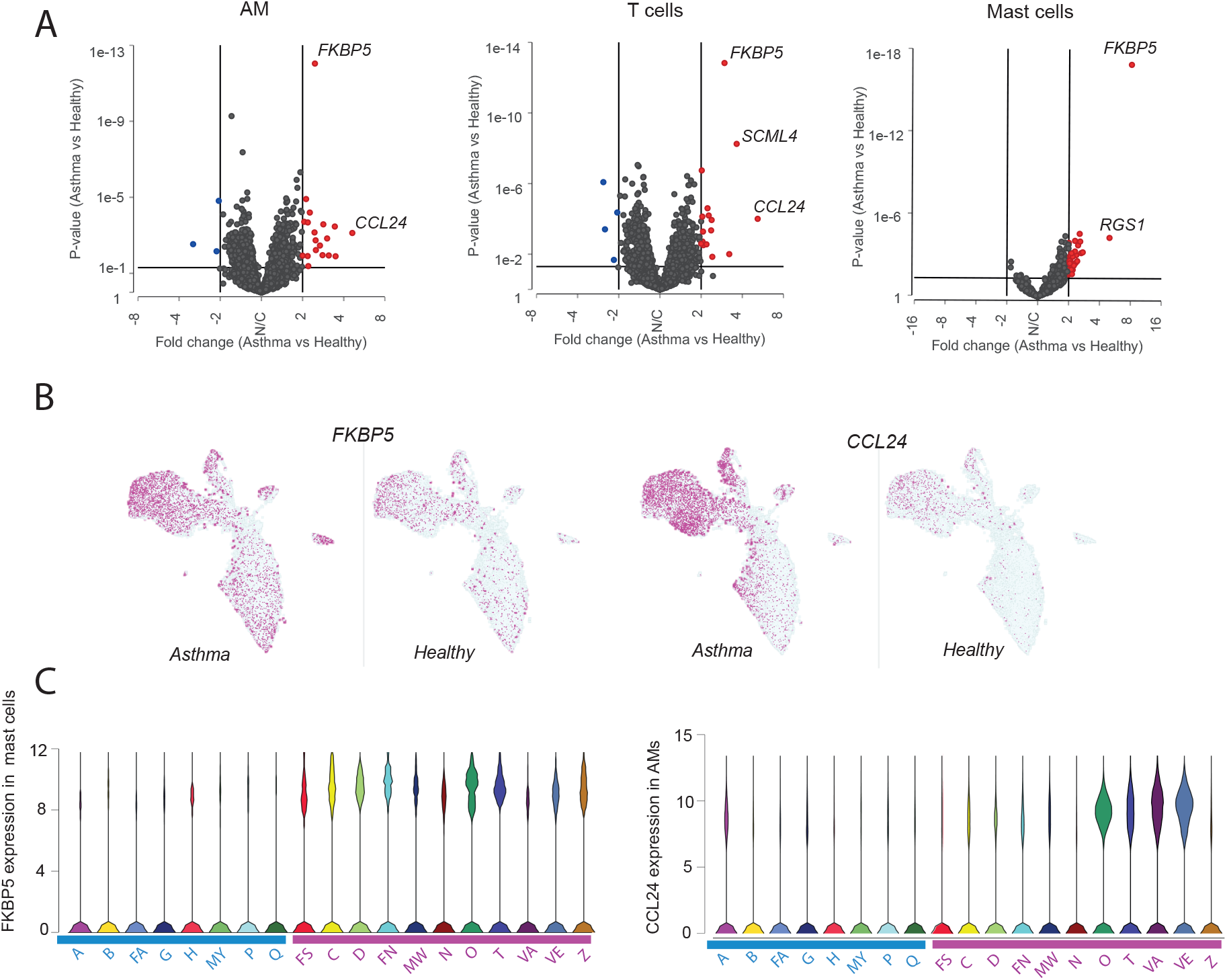
Differential gene expression EA and control horses. A) Volcano plots showing DE genes from pseudo-bulk (DESeq2) analysis of AMs, T-cells, and mast cells. B) UMAP visualization of *FKBP5* and *CCL25* expression in EA and control cells C) Violin plots showing expression levels of *FKBP5* in mast cells and *CCL24* in alveolar macrophages (AMs) for individual horses. IDs of healthy horses denoted in blue and asthma horses denoted in purple.

Several other significant DE genes were identified that have previously been associated with asthma in human or mice. Most of these genes were however detected as significant by only one or the other of the DE approaches (significance threshold = FDR < 0.1, FC ≥ 2, Supplementary File 5,6,7 and Supplementary Table 1). For instance, mast cells from asthma horses had higher expression of regulators of G-protein signaling *RGS1* and *RGS13*. Higher expression of thioredoxin interacting protein (*TXNIP*) were observed in mast cells as well as in AMs. Perilipin2 (*PLIN2*) and phospholipid-transporting ATPase (*ATP10A*) were upregulated in AM clusters AM4 and AM5, respectively (Supplementary Figure 7, Supplementary File 5 and 7).

When using MAST for DE between groups of neutrophils, several genes associated with inflammation were found to be significantly upregulated in the asthma horse cells, for instance *PGLYRP1, Eqca-1* (MHC class I heavy chain), *TXNIP, RSG2* and *IL4R* (Supplementary Table 1, Supplementary File 7). However, no significant gene hits came out in the neutrophil cluster when DE analysis was performed with the pseudo bulk (DESeq2) approach.

## Discussion

Mild-to moderate EA is one of the most common health and welfare issue in horses and inhibits performance in sport horses. Although EA has been studied extensively (12,39), little is known about the underlying mechanisms and the exact cellular phenotypes involved in the disease. mEA characterized by elevated mast cells in BAL and airway hypersensitiveness is a common type of asthma diagnosed at University Animal hospital in Sweden and all mEA cases included in the study were randomly selected clinical cases from the referral hospital.

Herein we report the first comprehensive scRNA-seq atlas of the immune cell populations from 63,022 cells in equine BAL sampled from 19 clinically examined horses with symptoms of mEA and healthy controls. Our study provides as high-resolution map of the cellular composition of BAL in health and mEA, as well as identified the glucocorticoid receptor associated protein FKBP5 as a biomarker candidate for diagnosis of EA.

Based on gene expression signatures we identified and categorized the major cell types present in equine BAL, namely; AMs, T cells, DCs, neutrophils and mast cells. The two largest equine BAL cell populations, which constituted > 90 % of cells in our data set, were T cells and AMs. We identified a diverse set of lung resident T cell subtypes, including CD4+ helper cells, CD8+ cytotoxic cells and a significant population of lung resident γδT cells, which are previously poorly characterized in horses (40). Interestingly, we identified inflammatory subsets of T cells (denoted T-ISG^Hi^) that exhibited strong interferon stimulation signatures. T-ISG^Hi^ cells were found among both the CD4+ and CD8+ tissue resident T cell groups. Although T-ISG^Hi^ cells were found in all horses, the strongest overall expressions of ISGs were found in a subset of the EA horses, suggesting either a specific asthma phenotype (Th1-driven) or a recent viral infection in those horses. This result warrants future studies with larger number of samples to determine if these T cell subsets have any functional implications for mEA

Human AMs have been shown to express high levels of *HLA-DR, CD11b (ITGAM), CD206 (MRC1), CD169 (SIGLEC1*) and *MARCO* (41). Likewise, we detected high levels of *HLA-DR, CD206* and *MARCO* transcripts in horse AMs, as well as moderate expression of *CD11b* and *CD169*. Flow cytometry experiments with human AMs have shown that expression of the scavenger receptor CD163 characterizes two subsets of AMs, and that CD163^Hi^ cells were more abundant than the CD163^Lo^ cells in BAL (41,42). Interestingly, we observed this pattern also in equine AMs. Our study revealed CD163 expression to be high in the majority of the equine AM populations whilst one cluster (denoted AM4) exhibited lower CD163 expression. Moreover, the AM4 cluster was transcriptionally very different from the other AMs, including lower levels of *MARCO*. Both human blood monocytes and mouse peritoneal macrophages have recently been shown to express very low levels of *MARCO* (43,44).

Although, embryonically derived AMs are known to be capable of self-proliferation (we did indeed observe a substantial number of proliferating AMs also in our horse data), it is also well established that the AM pool can be replenished by transformation of monocyte-derived macrophages. The recruitment of new AMs from monocyte precursors is frequently observed during e.g., infection and inflammation (41,45). The cells in the AM4 cluster expressed high levels of glycoprotein Nmb (*GPNMB*) as well as galectins (*LGALS1, LGALS3*), genes which have been shown to be upregulated during macrophage differentiation and in pro-inflammatory response (46,47). We therefore propose this AM cluster to comprise of recruited AMs and that those exhibit a more pro-inflammatory phenotype than the original population of resident AMs. This study did, however, not find significantly higher numbers of recruited AMs in the horses diagnosed with mEA.

Previous studies have reported significant amounts of functional T cell/monocyte complexes in blood which were found to mark specific immune perturbations (48). In this study we found a substantial number of cells with expression of both T cell and AM markers (double positive, DP cells). These DP cells presumably constitute doublets, either technical due to a probability of two cells entering in the same droplet, or they represent cell-cell complexes. To our knowledge there are no computational methods available to distinguish technical doublets from functional cell-cell complexes. As the number of cells clustering as DP in our analysis were rather high, we are tempted to speculate that a fraction of the DP cells constitutes functional T cell/AM complexes. Immune cells crosstalk is an essential feature of their function, being either performed by signaling molecules or by cells in functional complexes with each other (49). Nevertheless, macrophages are highly adherent cells and in future scRNA-seq studies involving equine BAL samples it may be worthwhile to consider additional strategies in order to ensure dissociation of cell-cell complexes prior to droplet encapsulation.

We also analyzed dendritic and granulocyte cells in BAL. Several functional classes of dendritic cells are found in humans, and some of those have previously been investigated in single-cell studies of BAL and other lung samples (36,50). In support of our annotations of the equine lung dendritic cell populations we found that the top markers identified for the different subclasses of dendritic cells in our study agree well with those identified for the human equivalents in single cell studies of the lung (36). Mast cells in equine BAL were found to be exclusively of the mucosal subtype. Several distinct subpopulations of neutrophils were identified, indicating high heterogeneity among the neutrophils infiltrating the equine airways. Given that rather small populations of these cell types were recapitulated and analyzed herein, further studies will be necessary to investigate the functional implications and impact of the identified subpopulations on equine airway disease

Interestingly, we identified highly significantly DE gene transcripts in clinical cases of mEA compared to healthy controls, which potentially can be further explored as potential biomarker for asthma and therapy response in horse and man. *FKBP5* was one of the most significant DE genes in all cell types analyzed (AMs, T cells and mast cells), and the highest upregulation was observed in the mast cells (Supplementary File 7). Although several studies have examined gene expression differences between healthy and asthmatic horses, to our knowledge there are no previous reports demonstrating any associations between *FKBP5* and EAS (22,51). The FKBP5 protein acts as a co-chaperone and modulates the glucocorticoid receptor activity by binding to other co-chaperones, e.g., the heat shock protein 90 (Hsp90) and P23 protein, which are important for the function of the glucocorticoid receptor (52,53). Hsp90 is one of the two main chaperone machines that influences glucocorticoid receptor assembly and activity, together with Hsp70 (54). By interacting with this complex, FKBP5 can modulate sensitivity of the glucocorticoid receptor. It has been demonstrated that increasing levels of FKBP5 reduces the transcriptional activity of the glucocorticoid receptor and that the binding of FKBP5 to Hsp90 affects cortisol affinity to the receptor (55). In squirrel monkeys, it has been reported that corticosteroid resistance is linked to overexpression of FKBP5 (56–58). In humans, about 10-20 % of patients with inflammatory diseases and immune disorders do not respond to treatment with glucocorticoids (59). In addition, corticosteroid insensitivity (i.e., an impaired response to corticosteroids) in asthma in humans is a well-recognized phenomenon, and it has been estimated that corticosteroid insensitivity is present in about one third of asthma patients (60). We are not aware of any published reports regarding the prevalence of corticosteroid resistance or insensitivity in horses. Mastocytic mEA is associated with airway hyperreactivity and affected horses can have limited response to corticosteroid treatment. Mast cell stabilizing drugs are sometimes suggested as an alterative treatment option. Considering this, the increased expression of *FKBP5* transcripts in lung immune cells of mEA horses with a mast cell phenotype is an important finding that is potentially related to steroid insensitivity in equines. It should be mentioned however that several factors, including disease severity, concurrent disease, antigen exposure, as well as the compliance and administration techniques by the caregivers, may have an effect on treatment response (61). As such, further studies are needed to fully understand the role of FKBP5 in asthma as well as in corticosteroid response in horses.

A second significantly DE gene, particularly highly expressed in AMs from mEAS horses, was *CCL24*, also known as eotaxin-2. *CCL24* is a cytokine belonging to the CC chemokine family. It interacts with the C-C Motif Chemokine Receptor 3 (*CCR3*) and stimulates eosinophilic chemotaxis (62). Several studies have demonstrated significant associations between the *CCL24* gene and human asthma, using airway epithelial cells and sputum (62–66). However, in BAL samples, while the levels of eotaxin-2 mRNA were higher than in the airway epithelial cells, the levels did not differ between asthmatic patients and healthy controls (62). The protein levels for eotaxin-2 were actually lower in the severe asthma cases, compared the healthy controls, and the severe group did not different from the milder asthma groups (62). In horses, a couple of studies have suggested a possible impact of *CCL24* on sEA (21,67). While the *CCL24* gene may be important for lung-eosinophilia, the current study did not observe any correlations between *CCL24* expression and individual eosinophil levels (as measured by cytology). Neither were there any differences in expression of *CCL24* when comparing asthmatic horses with high respectively low numbers of eosinophils. The *CCL24* expression was primarily upregulated in cells derived from a subset of the horses.

ScRNA-seq experiments performed on non-model organisms faces several challenges. One of the hurdles is that the equine reference genome is currently not as well annotated as the human or mouse genomes. Moreover, cell type annotation in horse is a laborious manual process as reference sets of cell-type markers is not yet available for this species. Another limitation of this study is the rather low sensitivity of the Drop-Seq technology, which restrains the number of genes that can be detected per cell. Moreover, droplet-based technologies are not optimal for analyzing sensitive cells such as granulocytes. However, as the field evolves toward more sensitive methods, we can expect increased resolution of cell-types and cell states in future scRNA-seq studies of EA.

## Conclusion

This study demonstrates for the first time the unbiased characterization of immune cells populating the horse airway in health and disease by scRNA-seq. Moreover, the detailed description of the cell types in BAL fluid will serve as a useful reference for future studies of equine asthma. Similarities with the immune cell subsets also found in human BAL-fluid strengthens the assumption that equine asthma can serve as a useful model for the human condition. Importantly, this study discloses the glucocorticoid receptor associated protein FKBP5 as a potential biomarker for EA.

## Materials and Methods

### BAL sampling and cytology analysis

All horses received detailed clinical and respiratory examinations, airway endoscopy.. Healthy control horses showed no signs of airway inflammation as judged by endoscopy examination. BAL fluid was collected after intravenous premedication with sedative (detomidine and butorphanol tartrate. A local anaesthetic was instilled at the trachea before performing BAL (Carbocain^®^). The BAL sampling was performed with a blind tube with 3 × 100 ml sterile isotonic saline solution (at 37 °C). A 15 ml aliquot of BAL fluid was transferred to the Clinical Pathology Laboratory, University Animal Hospital, Swedish University of Agricultural Sciences, and processed within 4 hours. Total nucleated cell count was determined using Advia 2120 (Siemens Healthcare GmbH, Ashburn, Germany). For cytological evaluation of BAL samples, cytospin preparations were prepared using two different concentrations of BAL-fluid. The first was prepared by centrifuging a 10 ml aliquot at 500 x g for 5 min and resuspending the resulting cell pellet with 50 μl albumin solution (1 g bovine serum albumin and 0.002 g NaN_3_ dissolved in 10 ml of 0.9% NaCl). The uncentrifuged preparation was made by adding 50 μl of the albumin solution to a 200 μl aliquot of the original BAL-fluid. Cytospin preparations with 100 μl aliquots were prepared in cytocentrifuge cassettes (Thermo Scientific Cytospin 4 centrifuge, Thermo Fisher Scientific, Waltham, Massachusetts, US). Slides were stained with May-GrÜnewald-Giemsa and evaluated by a clinical pathologist. A 200 differential count was performed where cells were classified as macrophages, lymphocytes, neutrophils, mast cells or eosinophils, expressed as a percentage. Epithelial cells, which often appear in aggregates, and nonintact cells were excluded from the differential count.

### Preparation and sequencing of Drop-Seq libraries

Cells were processed within 4 h of the BAL sampling. Single cells were encapsulated together with microbeads (ChemGenes Corporation, Wilmington, MA, USA) in droplets on the Nadia microfluidic device (Dolomite Bio, Royston, UK), according to the Drop-Seq protocol (27), with some minor modifications. Briefly, microbeads were resuspended in cell-lysis buffer prior to droplet encapsulation. The concentrations of the cell suspensions were kept low (300 cells/ul), to avoid aggregation of macrophages. The cell suspension was counted with the Cellometer K2 (Nexcelom Bioscience, Lawrence, MA, USA). Magnetic stirrers were used in the sample compartment of the Nadia Dolomite microfluidic cartridge. The microfluidic cartridge was loaded with 250 μl of cell suspension (300 cells/ul) and 250 μl of bead suspension (600 beads/ul). After droplet encapsulation, the resulting emulsion was collected and broken by filtration through a 5 um Uberstrainer filter (PluriSelect, Leipzig, Germany). The suspension was added to the filter and then 45 ml 6xSSC buffer was passed through the filter with the help of a 50 ml syringe. The beads were then collected from the filter, resuspended in high salt 6xSSC buffer, spun down and washed with 5x Maxima RT buffer (Thermo Fisher Scientific, Waltham, MA, USA). The beads were resuspended in 200 μl reverse transcription (RT) reaction mix containing: 1x Maxima H Minus RT buffer, 4% Ficoll PM-400, 2.5μM TSO oligo, 1 mM dNTP, RNase Inhibitors (RNaseOut and SUPERaseIn, ThermoFisher), 2000 U Maxima H Minus Reverse Transcriptase (Thermo Fisher Scientific, Waltham, MA, USA). The RT mix was incubated 30 min at 20°C followed by 90 min at 42°C and gentle shaking to keep the beads in suspension. After RT, Exonuclease I was added, in order to digest excess oligos, and incubated at 37°C for 30 min. The beads were spun down (1000g, 1 min) and washed, first in 1 ml TE-SDS buffer, then in 1×2 ml TE-Tween buffer and finally resuspended in H20. Beads were counted, aliquoted in a 96-well plate (5000 beads/well) and 14 cycles of cDNA amplification was then carried out using Terra PCR polymerase (TakaraBio, Kusatsu, Japan) and SMART PCR oligo. Amplified cDNA was pooled and purified twice with 0.6x AMPure beads (BeckmanCoulter, Brea, CA, USA). The concentration of amplified cDNA was measured with Qubit and 1 ng was used as input for Nextera XT (Illumina, San Diego, CA, USA) library preparation using New P5 SMART PCR oligo and Nextera index oligos. Phusion polymerase (Thermo Fisher Scientific, Waltham, MA, USA) was used for library amplification (10 PCR cycles). The number of beads corresponding to ~6000 cells/sample were taken into the amplification and library preparation steps. Three Nextera libraries per sample were prepared and pooled. Sequencing libraries were quantified using qPCR and sequenced on the NovaSeq6000 instrument (Illumina, San Diego, CA, USA) using custom sequencing primer (Read1CustomSeqB) and aiming at a read depth corresponding to > 50,000 reads/cell (Supplementary File 8). Primer sequences are listed in Supplementary Table 2.

### Primary data analysis and filtering

Data analysis and visualizations were performed using the Partek Flow single cell analysis framework (Partek Inc, Chesterfield, MO, USA) The Drop-Seq toolkit (27) implemented in Partek Flow were used to i) trim reads, ii) perform alignment to the EquCab3.0 (Ensembl release 97) using STAR v2.5.3a, iii) deduplicate UMIs, iv) filter and quantify barcodes to transcriptome. The EmptyDrops (68) test was used to distinguish real cells and empty droplets, which resulted in a recovery of between 2,538-5,757 cells per horse and in total 72,000 cells. Cells with fewer than 200 detected genes or with > 5 % mitochondrial reads were excluded. Potential doublets were identified and removed based on UMI counts > 8,000. This resulted in removal of 4 % of the sequenced cells (~3,000 cells). After the stringent quality filtering 63,000 cells remained for downstream analysis and clustering, in which between 200 and 1,620 genes/cells were detected. Genes with zero expression in > 99.9 % of cells were filtered out to reduce noise.

### Cluster analysis

Normalization (log2(CPM+1)) and scaling was performed followed by dimensional reduction in PCA. The number of principal components used in downstream clustering was determined based on Scree-plots. Harmony was used for batch-correction and integration of data from all horses (69). Graph-based clustering and UMAP was performed to visualize and annotate cell types (28,70). When re-clustering cell types which comprised only a small number of total cells the Harmony integration resulted in large numbers of non-informative clusters. The Harmony integration step was therefore omitted when re-clustering mast cells, neutrophils and DCs. Cluster biomarkers were computed by comparing the gene expression in every cluster to the other clusters combined, using the ‘compute biomarker’ function (t-test) implemented in Partek Flow. Differential gene expression between clusters or groups of clusters was also analyzed using the two-step hurdle model implemented in Partek Flow, which is equivalent to the MAST (‘Model-based Analysis of Single-cell Transcriptomics’) method (37). Cell types were manually annotated based on the expression of previously known and canonical cell type markers from studies of immune cells in humans and other animal models. Additional information regarding analysis parameter settings is listed in Supplementary File 8. The Kruskal-Wallis test was performed in order to test if cell group sizes (in %) were significantly different in EA vs healthy controls (Supplementary File 9)

### Differential gene expression between conditions

DE analysis across the EA and healthy groups was performed using i) a pseudo bulk approach and ii) a method specifically adapted for analyzing single cell data (two-step hurdle model, MAST) (37) as implemented in the Partek Flow software framework. For pseudo bulk analysis the gene expression values were pooled for every cell type or cluster analyzed, and differential gene expression across groups were computed using DESeq2. P-values were corrected using FDR multiple test correction (71). Additional information regarding parameter settings is found in Supplementary File 8. GO enrichment analysis was performed with g:Profiler (72).

## Declarations

### Ethics approval and consent to participate

All aspects of the study that involve sampling of horses have been approved by the regional ethical review board (5.8.18-20690/2020). All horse owners approved the study by written consent.

### Consent for publication

Not applicable

### Availability of data and materials

All raw sequencing reads will be made available in the National Center for Biotechnology Information (NCBI) sequence read archive (SRA). under BioProject accession number xxxxxxxx.

### Author contributions

AR conceived and designed the study together with MR. AR supervised the study, performed experiments, data analysis and drafted the manuscript. MR performed clinical examinations, BAL sampling and provided the clinical expertise. KF analyzed data and performed experiments. JN advised on methods and data analysis. IW, SW, SA and LH provided expertise in immunology. All authors reviewed and edited the manuscript.

## Acknowledgements

Sequencing was performed by the SciLifeLab National Genomics Infrastructure in Uppsala (SNP&SEQ Technology Platform) funded by the Swedish Research Council. Computational resources were provided by the Swedish National Infrastructure for Computing (SNIC) also partially funded by the Swedish Research Council.

---

**Supplementary File 1** Major cell types cluster markers

**Supplementary File 2** T cell types cluster markers

**Supplementary File 3** Alveolar macrophage cluster markers

**Supplementary File 4** Mast cell, neutrophil & dendritic cell cluster markers

**Supplementary File 5** Lists of DE genes AMs asthma vs healthy

**Supplementary File 6** Lists of DE genes T cells asthma vs healthy

**Supplementary File 7** Lists of DE genes mast cells & neutrophils asthma vs healthy

**Supplementary File 8** Analysis parameter settings

**Supplementary File 9** Compositional analysis statistics

**Supplementary Figure 1.**
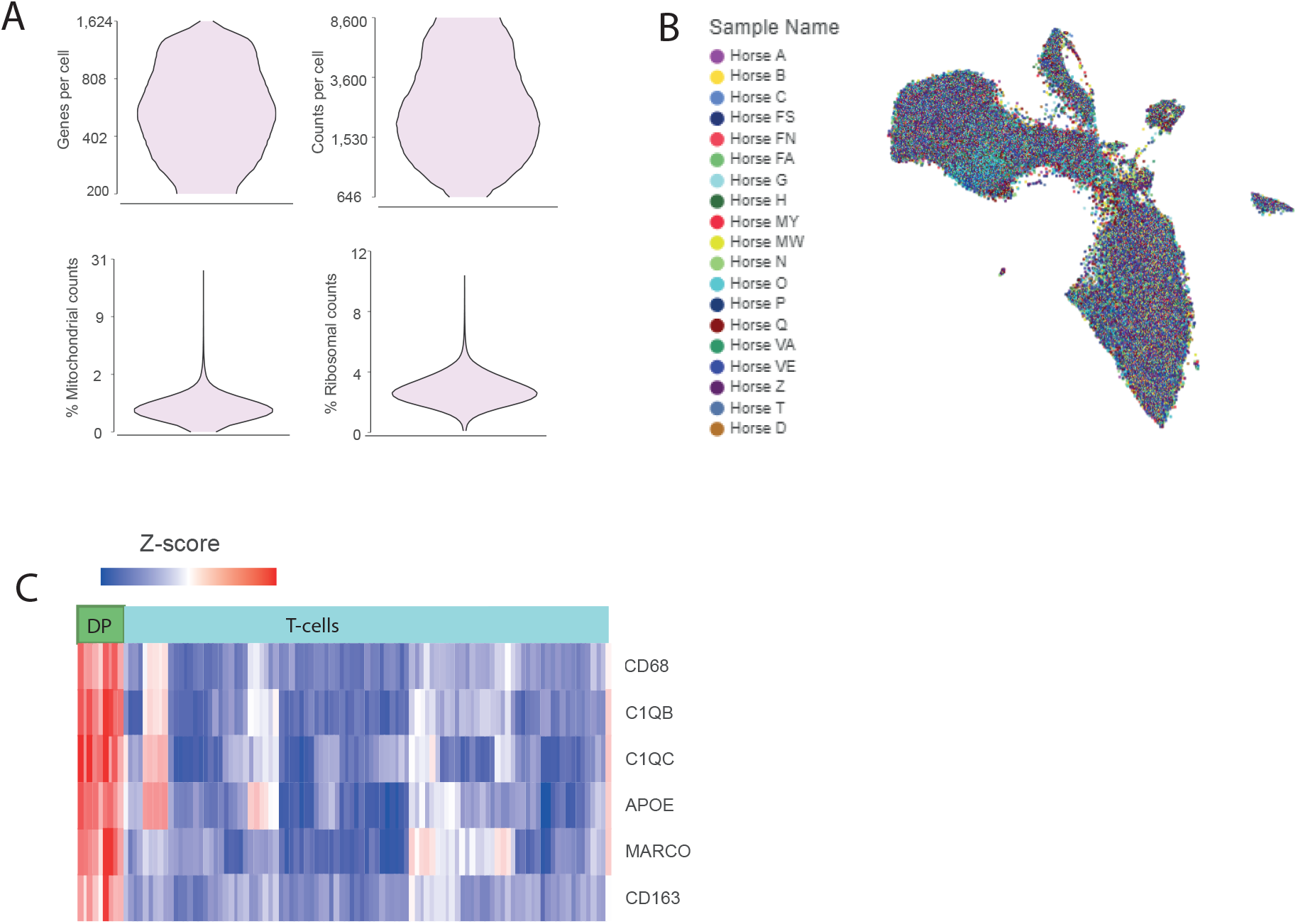
**A)** QC metrics shown for the 63,022 BAL cells after excluding low quality cells. B) UMAP visualization of integrated clustering of 63,022 BAL cells after batch cor-rection with Harmony. Cells are colored by sample name. C) Double positive (DP) cells clustered with the T-cells and at the same time demonstrated expression of macrophage markers. 2800 DP cells were excluded in downstream analysis.

**Supplementary Figure 2.**
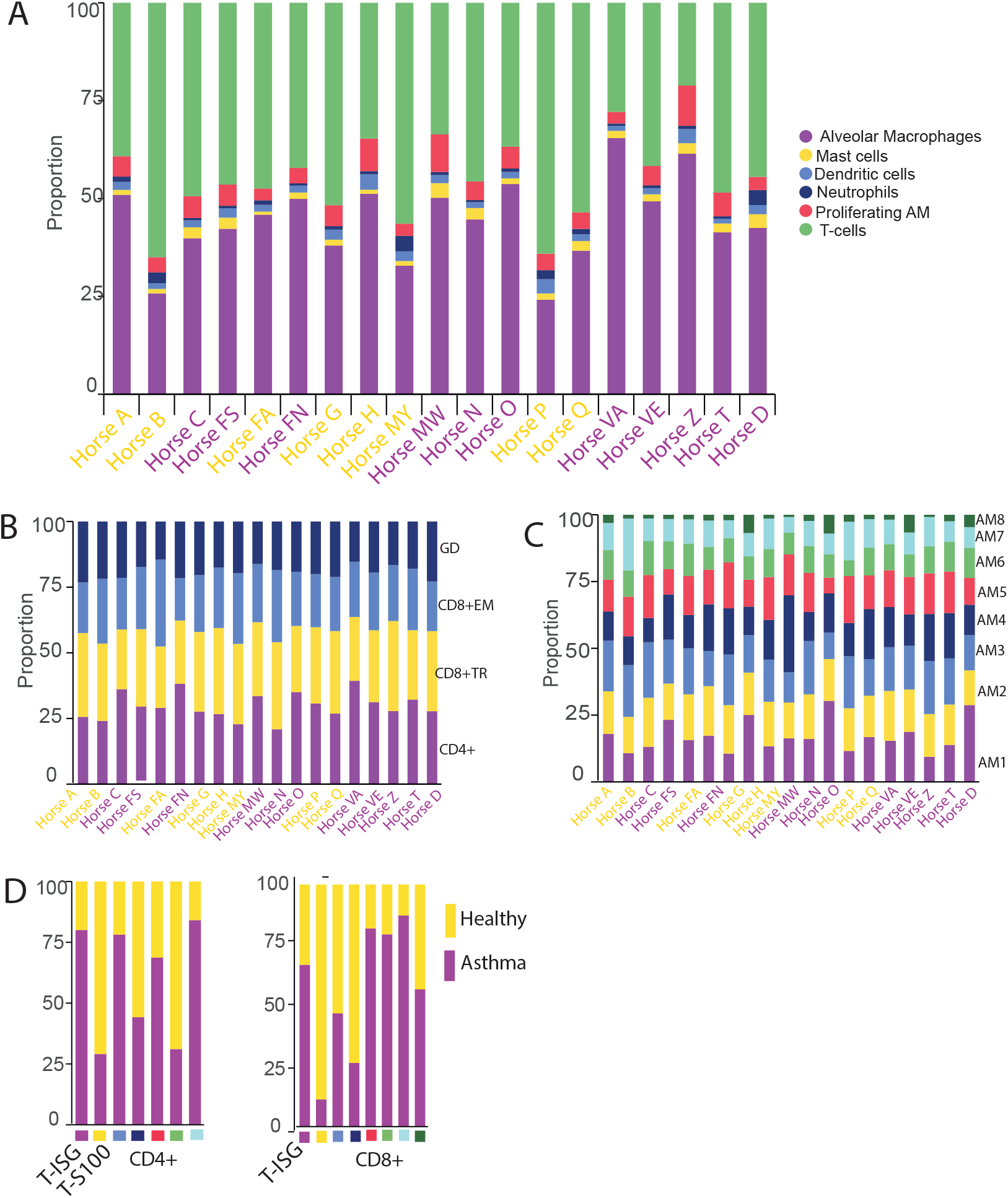
A) Proportions of cell types as identified by scRNA-seq in individual horses. B) Proportions of T cell subtypes in individual horses (GD=gamma delta T cells, EM=effector memory, TR = tissue resident). C) Proportions of AM clusters in individual horses D) Proportions of cells derived from asthma horses (purple bars) and healthy horses (yellow bars) respectively (T-ISG and T-S100) in CD4^+^ and CD8^+^ T cell iterative clustering. Colors below bars indicates the CD4^+^ and CD8^+^_TR_ sub-clusters and the color code is the same as in Figure 2D and E.

**Supplementary Figure 3.**
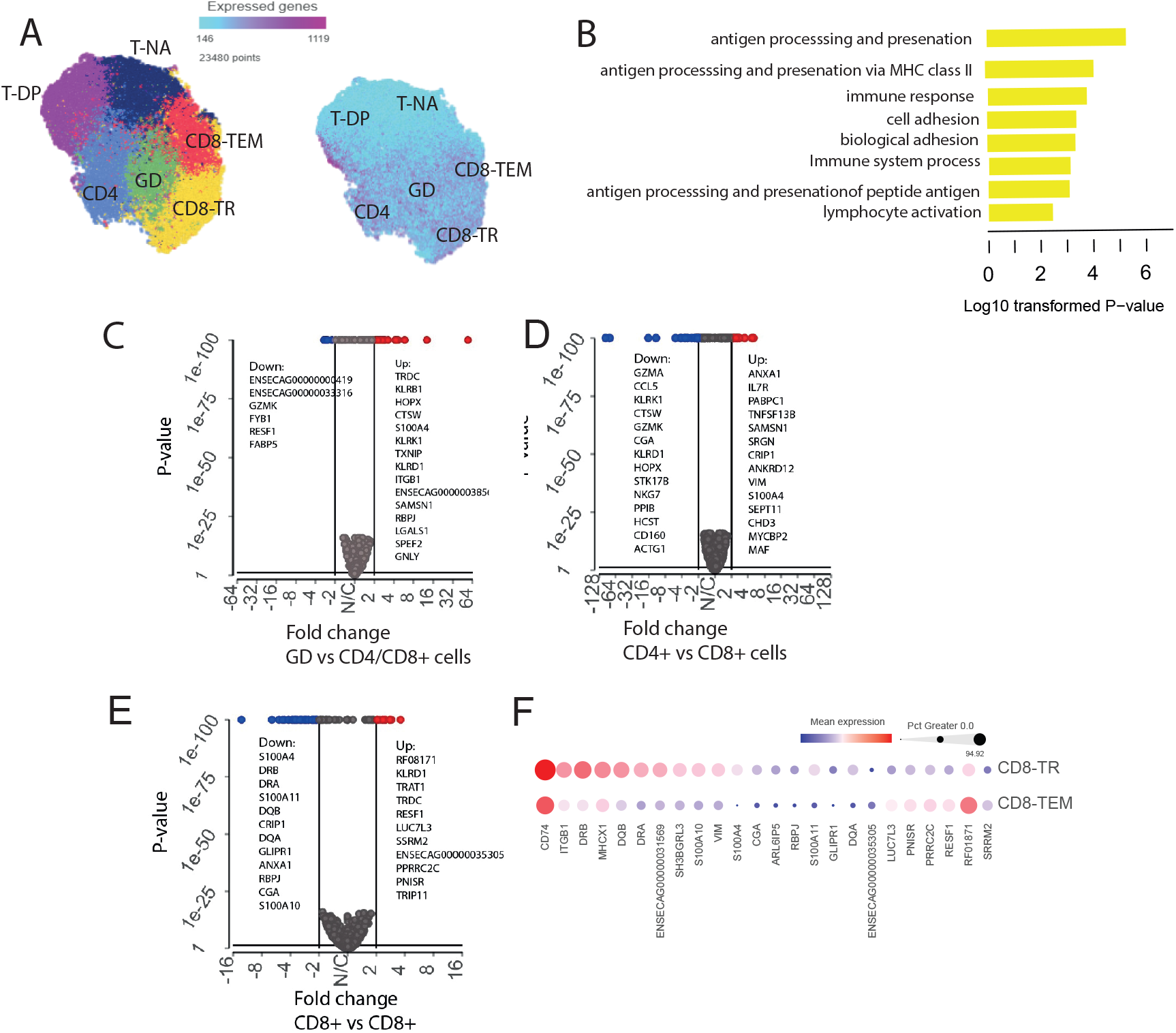
A) UMAP visualization of the six original T cell clusters (left) and the number of detected genes/cells mapped onto the UMAP plot (right). The cells in cluster T_NA_ were characterized by lower number of genes detected and displayed only subtle differential patterns (compared to the other T cell clusters, Supplementary File 2). Furthermore, the cluster denoted T_DP_ exhibited slightly increased expression of AM markers compared to the other T cell clusters. B) GO pathways upregulated in the CD8^+^TR cluster. C) Volcano plots showing DE analysis (MAST) of γδT cells compared to CD4^+^ and CD8^+^ cells. D) CD4^+^ vs CD8^+^ cells. E) CD8^+^_EM_ vs CD8^+^_TR_. F) Number of genes differentially expressed across the two CD8+ populations (FC>2) visualized in a bubble plot.

**Supplementary Figure 4.**
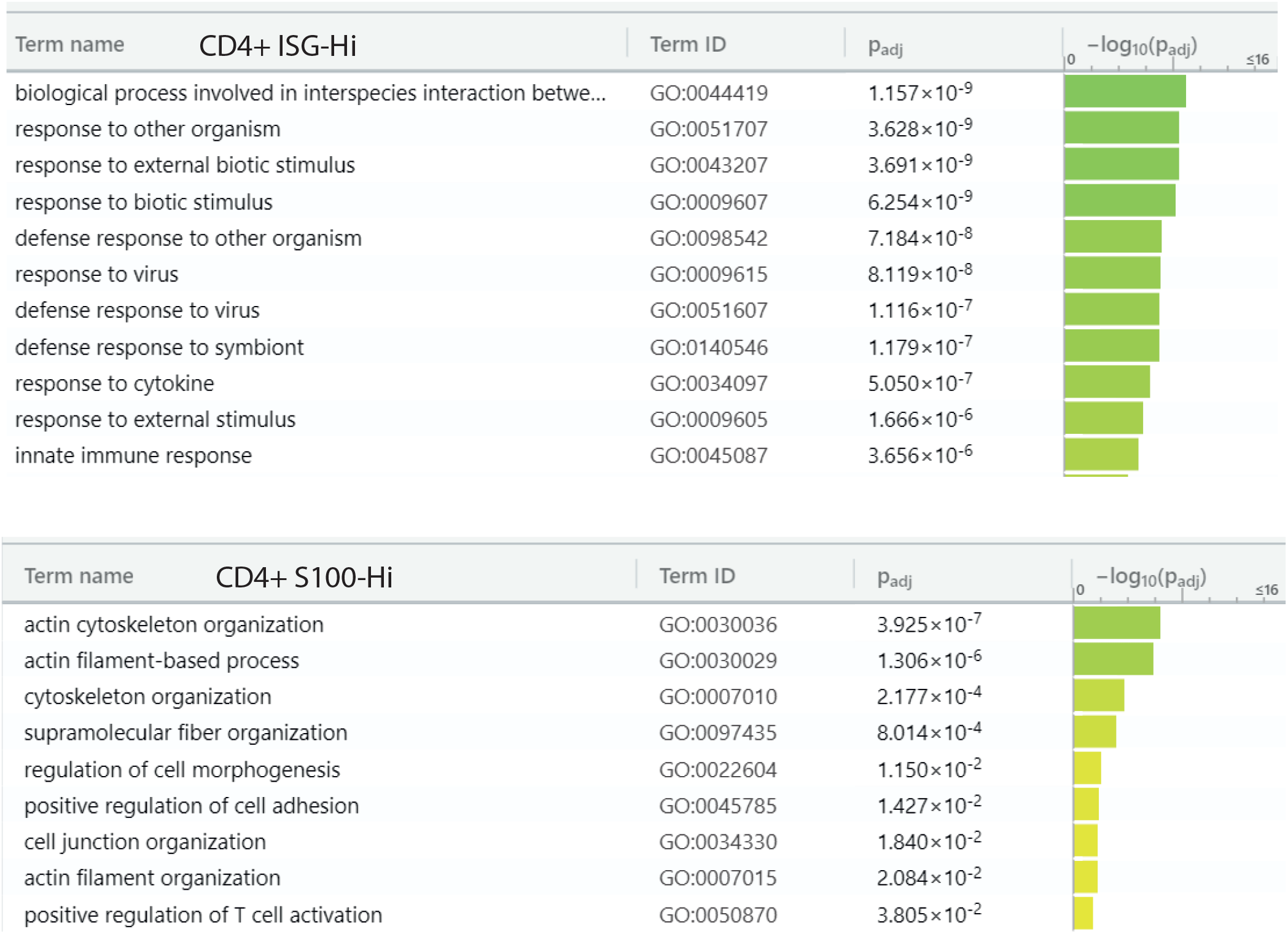
Upregulated GO pathways in CD4^+^ISG^Hi^ and CD4+ S100^Hi^ subclusters.

**Supplementary Figure 5.**
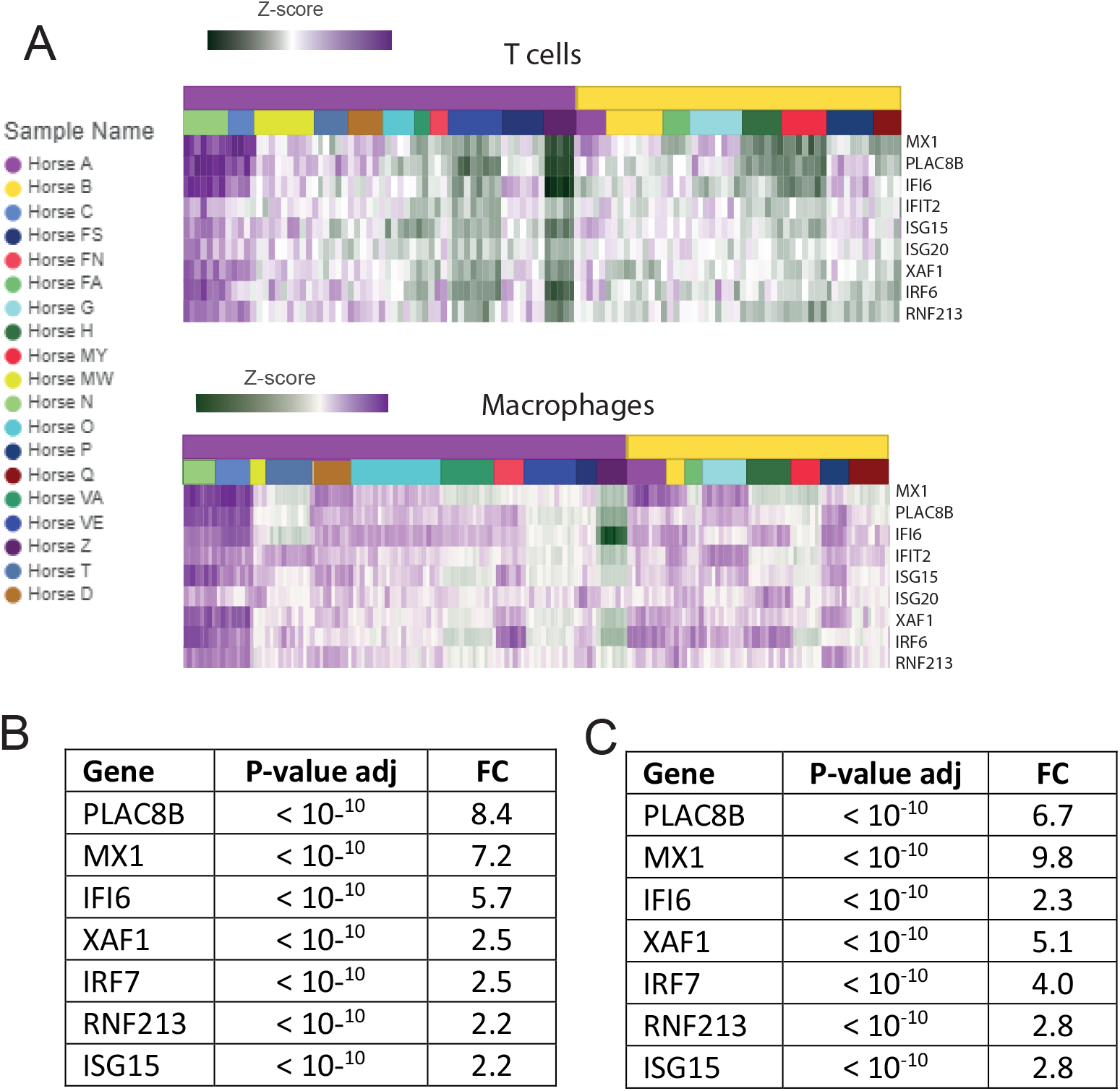
A) Heatmaps showing expression of a subset of interferon stimulated genes by individual horse and group (asthma=purple block, healthy= yellow block) in T cells and AMs, respectively. B) Differential expression (MAST) of a subset interferon stimulated in T cells. T cells from the two horses with the strongest ISG-signatures (Horse N and C) were tested against the T cells of the other EA horses. C) Differential expression (MAST) of a subset interferon stimulated in AMs. AMs from the two horses with the strongest ISG-signatures (Horse N and C) were tested against AMs of the other EA horses

**Supplementary Figure 6.**
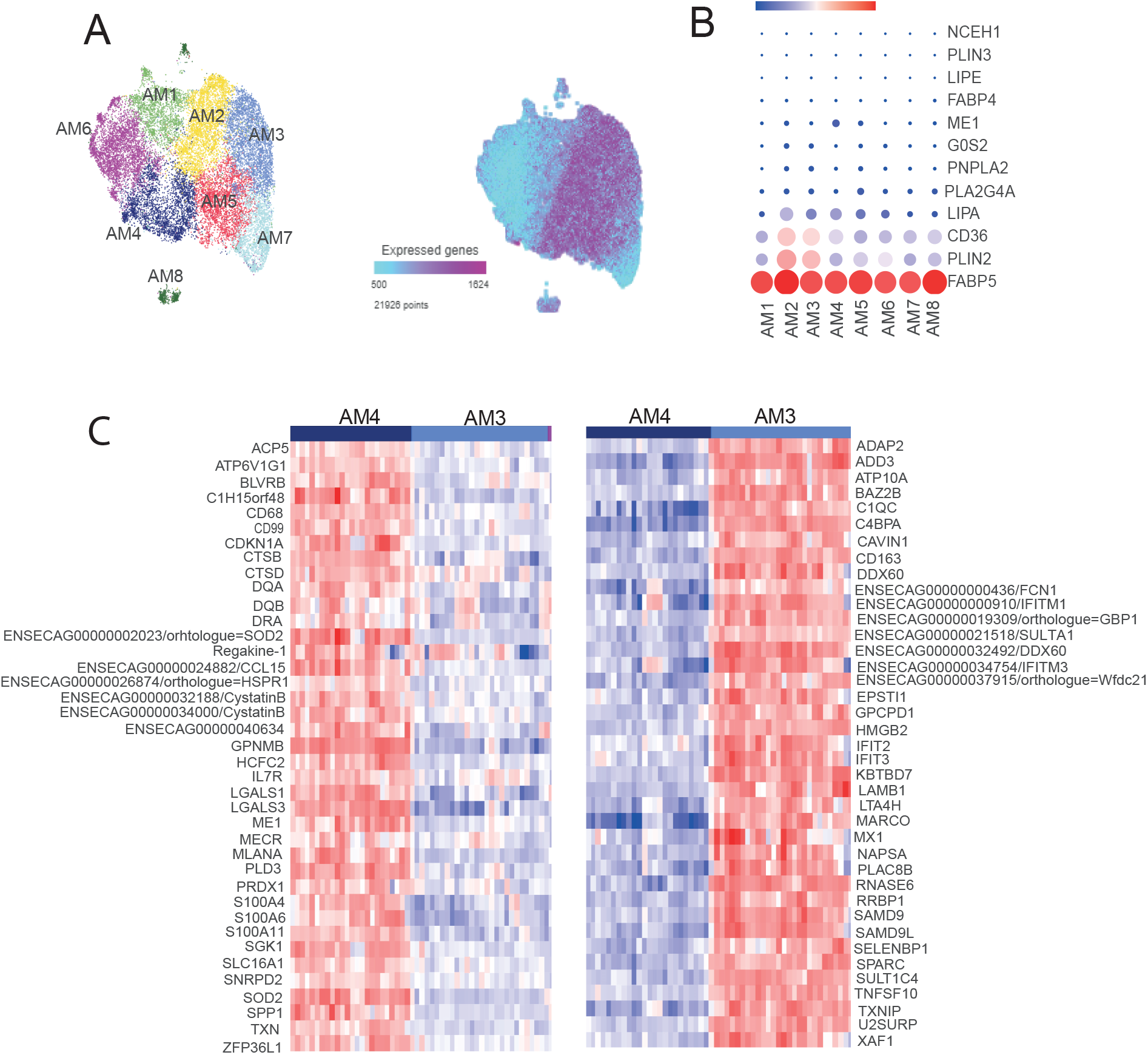
A) Number of detected genes/cells mapped onto the UMAP plot for AMs. B) Bubble plot showing expression of a subset of genes known to be involved in lipolysis and possibly M2 activation. C) Heatmaps showing genes top significantly upregulated (red) genes in AM clusters AM4 vs AM3 and vice versa.

**Supplementary Figure 7.**
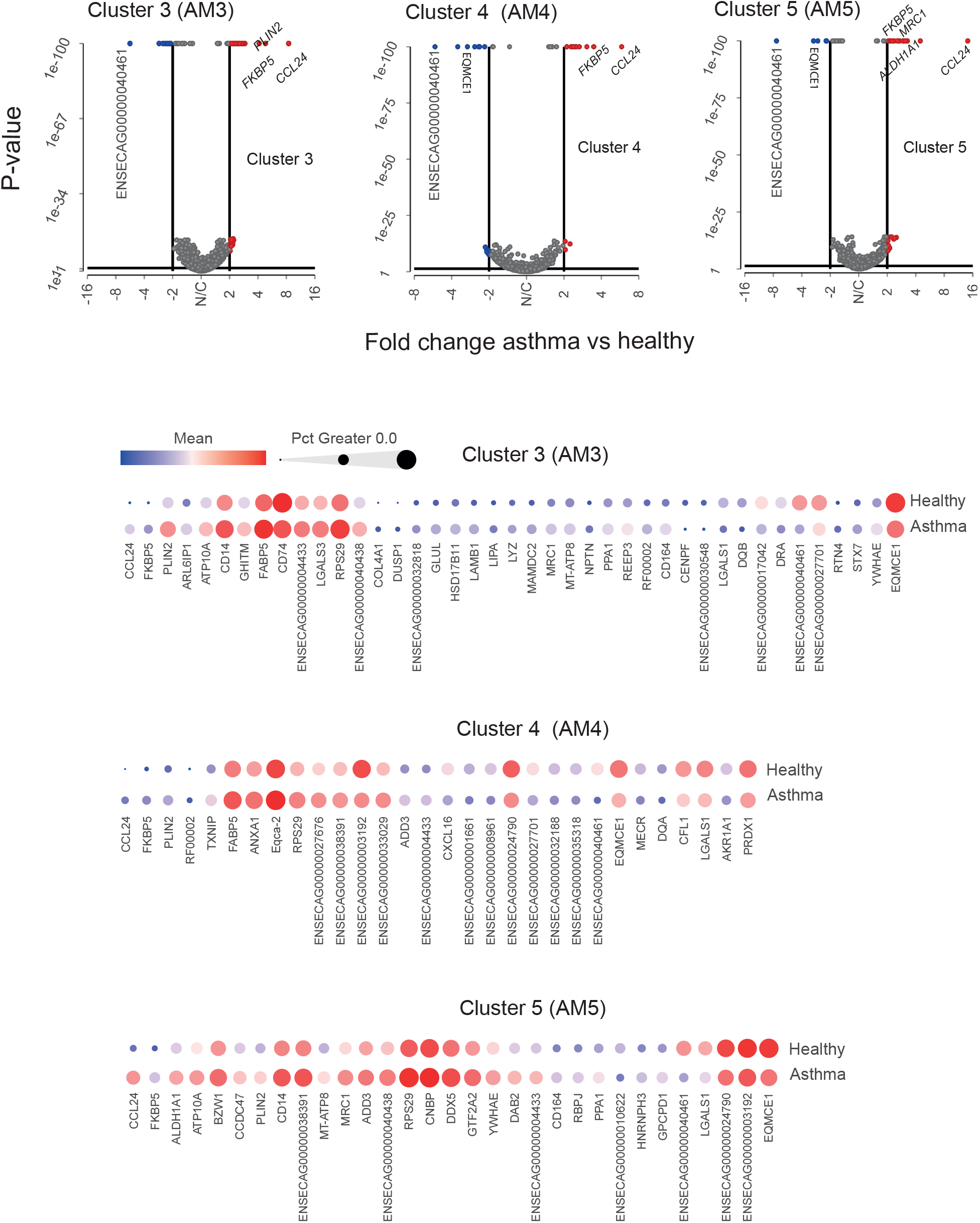
Differential expression analysis between asthma and healthy horses performed cluster-wise in alveolar macrophages subpopulations AM3, AM3, AM4. Differential expression visualized in volcano plots (top row) as well as in bubble plots.

**Supplementary Table 1.** Examples of DE genes (asthma vs healthy) See Supplementary Files 5,6,7 for complete lists. Only Hurdle model DE tests were performed on sub-cluster level (i.e alveolar macrophage clusters AM3, AM4, AM5)

| Genes up in asthma horses | Pseudo bulk DE<br>FC >2, FDR <0.01 | Single cell DE<br>FC >2, FDR <0.01 |
| --- | --- | --- |
| FKBP5 | AM, T-cells, Mast cells | AM, T-cells, Mast cells |
| CCL24 | AM, T-cells, | AM |
| RGS1 | mast cells | mast cells |
| RGS13 |  | mast cells |
| TXNIP |  | MA4, mast cells, neutrophils |
| PLIN2 | n.d | AM |
| ATP10A | n.d | AM |
| PECAM1 | AM | n.d |
| ENSECAG00000034004/O=FCGER1A | AM | n.d |
| GOS2 | AM | n.d |
| DUSP1 | AM | neutrophils |
| CD164 |  | MA3, MA4 |
| GLUL | n.d | AM |
| LAMB1 |  | MA3 |
| ANXA1 |  | MA4 |
| MRC1 |  | MA5 |
| ALDH1A1 |  | MA5 |
| ENSECAG00000038391/Eqca-1 | mast cells | T-cells, mast cells, neutrophils, MA5 |
| ENSECAG00000036180/novel gene | mast cells | mast cells |
| EPB41L2 | n.d | mast cells |
| ENSECAG00000032321/O=PGLYRP1 | n.d | neutrophils |
| IL4R | n.d | neutrophils |

n.d = not detected (or not significant, FC<2)

**Supplementary Table 2.**
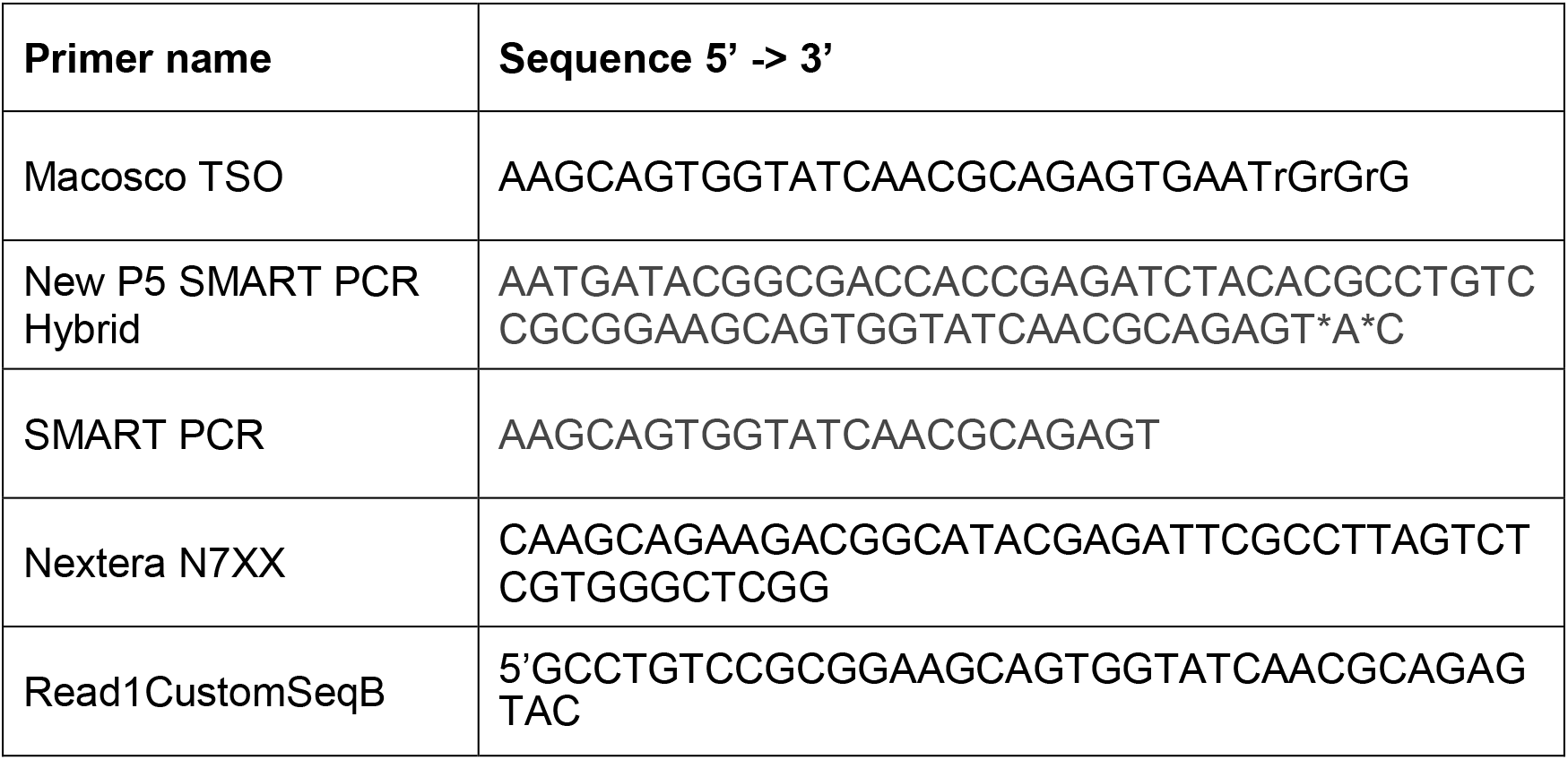
Primer Sequences

## References

1. Aun M, Bonamichi-Santos R, Arantes-Costa FM, Kalil J, Giavina-Bianchi P. Animal models of asthma: utility and limitations. J Asthma Allergy. 2017 Nov;Volume10:293–301.

2. Mullane K, Williams M. Animal models of asthma: Reprise or reboot? Biochem Pharmacol. 2014 Jan;87(1):131–9.

3. Holcombe SJ, Jackson C, Gerber V, Jefcoat A, Berney C, Eberhardt S, et al. Stabling is associated with airway inflammation in young Arabian horses. Equine Vet J. 2010 Jan 5;33(3):244–9.

4. Ramseyer A, Gaillard C, Burger D, Straub R, Jost U, Boog C, et al. Effects of Genetic and Environmental Factors on Chronic Lower Airway Disease in Horses. J Vet Intern Med. 2007 Jan;21(1):149–56.

5. Jost U, Klukowska-Rötzler J, Dolf G, Swinburne JE, Ramseyer A, Bugno M, et al. A region on equine chromosome 13 is linked to recurrent airway obstruction in horses. Equine Vet J. 2007 May;39(3):236–41.

6. Gerber V, Tessier C, Marti E. Genetics of upper and lower airway diseases in the horse: Genetics of upper and lower airway diseases in the horse. Equine Vet J. 2015 Jul;47(4):390–7.

7. Pacholewska A, Kraft M, Gerber V, Jagannathan V. Differential Expression of Serum MicroRNAs Supports CD4+ T Cell Differentiation into Th2/Th17 Cells in Severe Equine Asthma. Genes. 2017 Dec 12;8(12):383.

8. Mason VC, Schaefer RJ, McCue ME, Leeb T, Gerber V. eQTL discovery and their association with severe equine asthma in European Warmblood horses. BMC Genomics. 2018 Dec;19(1):581.

9. Gerber V. Genetics of Equine Respiratory Disease. Vet Clin North Am Equine Pract. 2020 Aug;36(2):243–53.

10. Gerber V, Baleri D, Klukowska-Rötzler J, Swinburne JE, Dolf G. Mixed Inheritance of Equine Recurrent Airway Obstruction. J Vet Intern Med. 2009 May;23(3):626–30.

11. Bond S, Léguillette R, Richard EA, Couetil L, Lavoie JP, Martin JG, et al. Equine asthma: Integrative biologic relevance of a recently proposed nomenclature. J Vet Intern Med. 2018 Nov;32(6):2088–98.

12. Couetil L, Cardwell JM, Leguillette R, Mazan M, Richard E, Bienzle D, et al. Equine Asthma: Current Understanding and Future Directions. Front Vet Sci. 2020 Jul 30;7:450.

13. Rettmer H, Hoffman AM, Lanz S, Oertly M, Gerber V. Owner-reported coughing and nasal discharge are associated with clinical findings, arterial oxygen tension, mucus score and bronchoprovocation in horses with recurrent airway obstruction in a field setting: Coughing and nasal discharge in equine recurrent airway obstruction. Equine Vet J. 2015 May;47(3):291–5.

14. Robinson NE, Berney C, Eberhart S, deFeijter-Rupp HeatherL, Jefcoat AM, Cornelisse CJ, et al. Coughing, mucus accumulation, airway obstruction, and airway inflammation in control horses and horses affected with recurrent airway obstruction. Am J Vet Res. 2003 May;64(5):550–7.

15. Bullone M, Lavoie JP. Asthma “of horses and men”—How can equine heaves help us better understand human asthma immunopathology and its functional consequences? Mol Immunol. 2015 Jul;66(1):97–105.

16. Couёtil LL, Cardwell JM, Gerber V, Lavoie J -P., Léguillette R, Richard EA. Inflammatory Airway Disease of Horses—Revised Consensus Statement. J Vet Intern Med. 2016 Mar;30(2):503–15.

17. Kuruvilla ME, Lee FEH, Lee GB. Understanding Asthma Phenotypes, Endotypes, and Mechanisms of Disease. Clin Rev Allergy Immunol. 2019 Apr;56(2):219–33.

18. Hulliger MF, Pacholewska A, Vargas A, Lavoie JP, Leeb T, Gerber V, et al. An Integrative miRNA-mRNA Expression Analysis Reveals Striking Transcriptomic Similarities between Severe Equine Asthma and Specific Asthma Endotypes in Humans. Genes. 2020 Sep 28;11(10):1143.

19. Cian F, Monti P, Durham A. Cytology of the lower respiratory tract in horses: An updated review. Equine Vet Educ. 2015 Oct;27(10):544–53.

20. Couetil LL, Thompson CA. Airway Diagnostics. Vet Clin North Am Equine Pract. 2020 Apr;36(1):87–103.

21. Pacholewska A, Jagannathan V, Drögemüller M, Klukowska-Rötzler J, Lanz S, Hamza E, et al. Impaired Cell Cycle Regulation in a Natural Equine Model of Asthma. Wade C, editor. PLOS ONE. 2015 Aug 20;10(8):e0136103.

22. Tessier L, Côté O, Clark ME, Viel L, Diaz-Méndez A, Anders S, et al. Impaired response of the bronchial epithelium to inflammation characterizes severe equine asthma. BMC Genomics. 2017 Dec;18(1):708.

23. Tessier L, Côté O, Bienzle D. Sequence variant analysis of RNA sequences in severe equine asthma. PeerJ. 2018 Oct 11;6:e5759.

24. Tessier L, Côté O, Clark ME, Viel L, Diaz-Méndez A, Anders S, et al. Gene set enrichment analysis of the bronchial epithelium implicates contribution of cell cycle and tissue repair processes in equine asthma. Sci Rep. 2018 Dec;8(1):16408.

25. Harman RM, Patel RS, Fan JC, Park JE, Rosenberg BR, Van de Walle GR. Single-cell RNA sequencing of equine mesenchymal stromal cells from primary donor-matched tissue sources reveals functional heterogeneity in immune modulation and cell motility. Stem Cell Res Ther. 2020 Dec;11(1):524.

26. Patel RS, Tomlinson JE, Divers TJ, Van de Walle GR, Rosenberg BR. Single-cell resolution landscape of equine peripheral blood mononuclear cells reveals diverse cell types including T-bet+ B cells. BMC Biol. 2021 Dec;19(1):13.

27. Macosko EZ, Basu A, Satija R, Nemesh J, Shekhar K, Goldman M, et al. Highly Parallel Genome-wide Expression Profiling of Individual Cells Using Nanoliter Droplets. Cell. 2015 May;161(5):1202–14.

28. McInnes L, Healy J, Saul N, Großberger L. UMAP: Uniform Manifold Approximation and Projection. J Open Source Softw. 2018 Sep 2;3(29):861.

29. Burel JG, Pomaznoy M, Lindestam Arlehamn CS, Seumois G, Vijayanand P, Sette A, et al. The Challenge of Distinguishing Cell–Cell Complexes from Singlet Cells in Non-Imaging Flow Cytometry and Single-Cell Sorting. Cytometry A. 2020 Nov;97(11):1127–35.

30. Burel JG, Pomaznoy M, Lindestam Arlehamn CS, Weiskopf D, da Silva Antunes R, Jung Y, et al. Circulating T cell-monocyte complexes are markers of immune perturbations. eLife. 2019 Jun 25;8:e46045.

31. Choi H, Song H, Jung YW. The Roles of CCR7 for the Homing of Memory CD8+ T Cells into Their Survival Niches. Immune Netw. 2020;20(3):e20.

32. Mould KJ, Jackson ND, Henson PM, Seibold M, Janssen WJ. Single cell RNA sequencing identifies unique inflammatory airspace macrophage subsets. JCI Insight. 2019 Mar 7;4(5):e126556.

33. Wang L, Netto KG, Zhou L, Liu X, Wang M, Zhang G, et al. Single-cell transcriptomic analysis reveals the immune landscape of lung in steroid-resistant asthma exacerbation. Proc Natl Acad Sci. 2021 Jan 12;118(2):e2005590118.

34. Gibbings SL, Goyal R, Desch AN, Leach SM, Prabagar M, Atif SM, et al. Transcriptome analysis highlights the conserved difference between embryonic and postnatal-derived alveolar macrophages. Blood. 2015 Sep 10;126(11):1357–66.

35. Irani AA, Schechter NM, Craig SS, DeBlois G, Schwartz LB. Two types of human mast cells that have distinct neutral protease compositions. Proc Natl Acad Sci. 1986 Jun;83(12):4464–8.

36. Naessens T, Morias Y, Hamrud E, Gehrmann U, Budida R, Mattsson J, et al. Human Lung Conventional Dendritic Cells Orchestrate Lymphoid Neogenesis during Chronic Obstructive Pulmonary Disease. Am J Respir Crit Care Med. 2020 Aug 15;202(4):535–48.

37. Finak G, McDavid A, Yajima M, Deng J, Gersuk V, Shalek AK, et al. MAST: a flexible statistical framework for assessing transcriptional changes and characterizing heterogeneity in single-cell RNA sequencing data. Genome Biol. 2015 Dec;16(1):278.

38. Love MI, Huber W, Anders S. Moderated estimation of fold change and dispersion for RNA-seq data with DESeq2. Genome Biol. 2014 Dec;15(12):550.

39. Simões J, Batista M, Tilley P. The Immune Mechanisms of Severe Equine Asthma—Current Understanding and What Is Missing. Animals. 2022 Mar 16;12(6):744.

40. Tallmadge RL, Wang M, Sun Q, Felippe MJB. Transcriptome analysis of immune genes in peripheral blood mononuclear cells of young foals and adult horses. Bielefeldt-Ohmann H, editor. PLOS ONE. 2018 Sep 5;13(9):e0202646.

41. Bain CC, MacDonald AS. The impact of the lung environment on macrophage development, activation and function: diversity in the face of adversity. Mucosal Immunol. 2022 Feb;15(2):223–34.

42. Bharat A, Bhorade SM, Morales-Nebreda L, McQuattie-Pimentel AC, Soberanes S, Ridge K, et al. Flow Cytometry Reveals Similarities Between Lung Macrophages in Humans and Mice. Am J Respir Cell Mol Biol. 2016 Jan;54(1):147–9.

43. Lara S, Akula S, Fu Z, Olsson AK, Kleinau S, Hellman L. The Human Monocyte—A Circulating Sensor of Infection and a Potent and Rapid Inducer of Inflammation. Int J Mol Sci. 2022 Mar 31;23(7):3890.

44. Paivandy A, Akula S, Lara S, Fu Z, Olsson AK, Kleinau S, et al. Quantitative In-Depth Transcriptome Analysis Implicates Peritoneal Macrophages as Important Players in the Complement and Coagulation Systems. Int J Mol Sci. 2022 Jan 21;23(3):1185.

45. Evren E, Ringqvist E, Willinger T. Origin and ontogeny of lung macrophages: from mice to humans. Immunology. 2020 Jun;160(2):126–38.

46. Ripoll VM, Irvine KM, Ravasi T, Sweet MJ, Hume DA. *Gpnmb* Is Induced in Macrophages by IFN-γ and Lipopolysaccharide and Acts as a Feedback Regulator of Proinflammatory Responses. J Immunol. 2007 May 15;178(10):6557–66.

47. Yaseen H, Butenko S, Polishuk-Zotkin I, Schif-Zuck S, Pérez-Sáez JM, Rabinovich GA, et al. Galectin-1 Facilitates Macrophage Reprogramming and Resolution of Inflammation Through IFN-β. Front Pharmacol. 2020 Jun 17;11:901.

48. Bloom JD. Estimating the frequency of multiplets in single-cell RNA sequencing from cell-mixing experiments. PeerJ. 2018 Sep 3;6:e5578.

49. Guerriero JL. Macrophages. In: International Review of Cell and Molecular Biology [Internet]. Elsevier; 2019 [cited 2022 Sep 13]. p. 73–93. Available from: https://linkinghub.elsevier.com/retrieve/pii/S1937644818300698

50. Aegerter H, Lambrecht BN, Jakubzick CV. Biology of lung macrophages in health and disease. Immunity. 2022 Sep;55(9):1564–80.

51. Davis KU, Sheats MK. Differential gene expression and Ingenuity Pathway Analysis of bronchoalveolar lavage cells from horses with mild/moderate neutrophilic or mastocytic inflammation on BAL cytology. Vet Immunol Immunopathol. 2021 Apr;234:110195.

52. Schiene-Fischer C, Yu C. Receptor accessory folding helper enzymes: the functional role of peptidyl prolyl *cis* / *trans* isomerases. FEBS Lett. 2001 Apr 20;495(1–2):1–6.

53. Kirschke E, Goswami D, Southworth D, Griffin PR, Agard DA. Glucocorticoid Receptor Function Regulated by Coordinated Action of the Hsp90 and Hsp70 Chaperone Cycles. Cell. 2014 Jun;157(7):1685–97.

54. Grad I, Picard D. The glucocorticoid responses are shaped by molecular chaperones. Mol Cell Endocrinol. 2007 Sep;275(1–2):2–12.

55. Wochnik GM, Rüegg J, Abel GA, Schmidt U, Holsboer F, Rein T. FK506-binding Proteins 51 and 52 Differentially Regulate Dynein Interaction and Nuclear Translocation of the Glucocorticoid Receptor in Mammalian Cells. J Biol Chem. 2005 Feb;280(6):4609–16.

56. Westberry JM, Sadosky PW, Hubler TR, Gross KL, Scammell JG. Glucocorticoid resistance in squirrel monkeys results from a combination of a transcriptionally incompetent glucocorticoid receptor and overexpression of the glucocorticoid receptor co-chaperone FKBP51. J Steroid Biochem Mol Biol. 2006 Jul;100(1–3):34–41.

57. Denny WB, Valentine DL, Reynolds PD, Smith DF, Scammell JG. Squirrel Monkey Immunophilin FKBP51 Is a Potent Inhibitor of Glucocorticoid Receptor Binding ^1^. Endocrinology. 2000 Nov;141(11):4107–13.

58. Scammell JG, Denny WB, Valentine DL, Smith DF. Overexpression of the FK506-Binding Immunophilin FKBP51 Is the Common Cause of Glucocorticoid Resistance in Three New World Primates. Gen Comp Endocrinol. 2001 Nov;124(2):152–65.

59. Panda L, Mabalirajan U. Recent Updates on Corticosteroid Resistance in Asthma. Emerg Med J. 2018;3(3):49–57.

60. Thomson NC. Addressing corticosteroid insensitivity in adults with asthma. Expert Rev Respir Med. 2016 Feb;10(2):137–56.

61. Mainguy-Seers S, Lavoie J. Glucocorticoid treatment in horses with asthma: A narrative review. J Vet Intern Med. 2021 Jul;35(4):2045–57.

62. Coleman JM, Naik C, Holguin F, Ray A, Ray P, Trudeau JB, et al. Epithelial eotaxin-2 and eotaxin-3 expression: relation to asthma severity, luminal eosinophilia and age at onset. Thorax. 2012 Dec;67(12):1061–6.

63. Ravensberg AJ, Ricciardolo FLM, van Schadewijk A, Rabe KF, Sterk PJ, Hiemstra PS, et al. Eotaxin-2 and eotaxin-3 expression is associated with persistent eosinophilic bronchial inflammation in patients with asthma after allergen challenge. J Allergy Clin Immunol. 2005 Apr;115(4):779–85.

64. Komiya A, Nagase H, Yamada H, Sekiya T, Yamaguchi M, Sano Y, et al. Concerted expression of eotaxin-1, eotaxin-2, and eotaxin-3 in human bronchial epithelial cells. Cell Immunol. 2003 Oct;225(2):91–100.

65. Berkman N, Ohnona S, Chung FK, Breuer R. Eotaxin-3 but Not Eotaxin Gene Expression Is Upregulated in Asthmatics 24 Hours after Allergen Challenge. Am J Respir Cell Mol Biol. 2001 Jun 1;24(6):682–7.

66. Scheicher ME, Teixeira MM, Cunha FQ, Teixeira AL, Filho JT, Vianna EO. Eotaxin-2 in sputum cell culture to evaluate asthma inflammation. Eur Respir J. 2007 Mar 1;29(3):489–95.

67. Swinburne JE, Bogle H, Klukowska-Rötzler J, Drögemüller M, Leeb T, Temperton E, et al. A whole-genome scan for recurrent airway obstruction in Warmblood sport horses indicates two positional candidate regions. Mamm Genome. 2009 Aug;20(8):504–15.

68. Lun ATL, Riesenfeld S, Andrews T, Dao TP, Gomes T, participants in the 1st Human Cell Atlas Jamboree, et al. EmptyDrops: distinguishing cells from empty droplets in droplet-based single-cell RNA sequencing data. Genome Biol. 2019 Dec;20(1):63.

69. Korsunsky I, Millard N, Fan J, Slowikowski K, Zhang F, Wei K, et al. Fast, sensitive and accurate integration of single-cell data with Harmony. Nat Methods. 2019 Dec;16(12):1289–96.

70. Becht E, McInnes L, Healy J, Dutertre CA, Kwok IWH, Ng LG, et al. Dimensionality reduction for visualizing single-cell data using UMAP. Nat Biotechnol. 2019 Jan;37(1):38–44.

71. Benjamini Y, Hochberg Y. Controlling the False Discovery Rate: A Practical and Powerful Approach to Multiple Testing. J R Stat Soc Ser B Methodol. 1995;57(1):289–300.

72. Raudvere U, Kolberg L, Kuzmin I, Arak T, Adler P, Peterson H, et al. g:Profiler: a web server for functional enrichment analysis and conversions of gene lists (2019 update). Nucleic Acids Res. 2019 Jul 2;47(W1):W191–8.

